# Longitudinal Multi-omic Immune Profiling Reveals Age-Related Immune Cell Dynamics in Healthy Adults

**DOI:** 10.1101/2024.09.10.612119

**Authors:** Qiuyu Gong, Mehul Sharma, Emma L. Kuan, Marla C. Glass, Aishwarya Chander, Mansi Singh, Lucas T. Graybuck, Zachary J. Thomson, Christian M. LaFrance, Samir Rachid Zaim, Tao Peng, Lauren Y. Okada, Palak C Genge, Katherine E. Henderson, Elisabeth M. Dornisch, Erik D. Layton, Peter J. Wittig, Alexander T. Heubeck, Nelson M. Mukuka, Julian Reading, Charles R. Roll, Veronica Hernandez, Vaishnavi Parthasarathy, Tyanna J. Stuckey, Blessing Musgrove, Elliott Swanson, Cara Lord, Morgan D.A. Weiss, Cole G. Phalen, Regina R. Mettey, Kevin J. Lee, John B. Johanneson, Erin K. Kawelo, Jessica Garber, Upaasana Krishnan, Megan Smithmyer, E. John Wherry, Laura Vella, Sarah E. Henrickson, Mackenzie S. Kopp, Adam K. Savage, Lynne A. Becker, Paul Meijer, Ernest M. Coffey, Jorg J. Goronzy, Cate Speake, Thomas F. Bumol, Ananda W. Goldrath, Troy R. Torgerson, Xiao-jun Li, Peter J. Skene, Jane H. Buckner, Claire E. Gustafson

## Abstract

The generation and maintenance of protective immunity is a dynamic interplay between host and environment that is impacted by age. Understanding fundamental changes in the healthy immune system that occur over a lifespan is critical in developing interventions for age-related susceptibility to infections and diseases. Here, we use multi-omic profiling (scRNA-seq, proteomics, flow cytometry) to examined human peripheral immunity in over 300 healthy adults, with 96 young and older adults followed over two years with yearly vaccination. The resulting resource includes scRNA-seq datasets of >16 million PBMCs, interrogating 71 immune cell subsets from our new Immune Health Atlas. This study allows unique insights into the composition and transcriptional state of immune cells at homeostasis, with vaccine perturbation, and across age. We find that T cells specifically accumulate age-related transcriptional changes more than other immune cells, independent from inflammation and chronic perturbation. Moreover, impaired memory B cell responses to vaccination are linked to a Th2-like state shift in older adults’ memory CD4 T cells, revealing possible mechanisms of immune dysregulation during healthy human aging. This extensive resource is provided with a suite of exploration tools at https://apps.allenimmunology.org/aifi/insights/dynamics-imm-health-age/ to enhance data accessibility and further the understanding of immune health across age.

## Introduction

Tracking the dynamics of the healthy immune landscape over the lifespan is critical for understanding susceptibility to infections, responses to vaccines, and the onset of immune-related diseases that occur differentially across the aging spectrum. While many studies on immune health and aging utilize single snapshots of the immune system to infer common features of this aging process, (Sayed et al. 2021; Whiting et al. 2015; Sparks et al. 2024) the function of immune cells is always dictated by some element of time. The innate immune compartment (i.e., monocytes, neutrophils) is heavily engaged in rapid and stochastic responses (hours to days) whereas the adaptive immune compartment (T cells, B cells) mediates slower but more durable memory responses (days to years). Indeed, recent studies focused on longitudinal monitoring in the context of infection, vaccination and homeostasis have offered a unique view of the global age-associated changes in the immune system and provided deeper insights into the dynamic interplay of immunity with exposures over time. (Fourati et al. 2022; Van Phan et al. 2024; Alpert et al. 2019)

The wide-spread implementation of single-cell RNA sequencing (scRNA-seq) has revolutionized our ability to dissect the complexities of the immune system, enabling deep interrogation of individual immune cells. (Terekhova et al. 2023; Mogilenko, Shchukina, and Artyomov 2022) Combining scRNA-seq, and other high-plex methods, with longitudinal sampling offers unprecedented insights into the ongoing adaptation of the immune system at a single cell level and interactions between immune cells and their surrounding microenvironment. Notably, memory T cells and B cells play pivotal roles in long-term immunity, collectively orchestrating responses to pathogens and vaccines throughout one’s life. Memory responses in mice and humans can be maintained for decades, in part through self-renewal. (Soerens et al. 2023; Akondy et al. 2017; Fuertes Marraco et al. 2015) However, in long-term mouse studies, memory T cells progressively accumulated unique transcriptional programs over time. (Soerens et al. 2023) Similarly, transcriptional alterations in the T cell compartment of humans have been increasingly recognized as a feature of aging (H. Zhang et al. 2022; Thomson et al. 2023; Moskowitz et al. 2017), associated with diminished immune responses and increased vulnerability to infections among older adults. (Gustafson et al. 2020) However, the breadth, variation and, in turn, stability of these changes across the entire peripheral immune compartment and their link to concurrent age-related immune dysfunction in people, including impaired vaccine-specific antibody production and a higher propensity for chronic viral re-activation, are not fully elucidated.

Here, we longitudinally profiled the peripheral immune system in 96 healthy young and older adults over 2 years, in the homeostatic state and following annual immune perturbation induced by influenza vaccination. Employing scRNA-seq, high-dimensional plasma proteomics, and spectral flow cytometry to samples collected at 8-10 time points per donor, we investigated the molecular and cellular mechanisms underlying broad age-related changes in immune responsiveness. This effort generated a human peripheral immune cell scRNA-seq reference dataset composed of over 13.7 million peripheral blood mononuclear cells (PBMCs) from our longitudinal, prospective cohort and an additional 3.2 million PBMCs from a cross-sectional follow-up cohort of 234 healthy adults. Our study uniquely demonstrates that T cell subsets in older adults maintain distinct transcriptional programming compared to those in young adults, with reprogramming in early T cell subsets accumulating over time, independent of inflammation and changes induced by chronic CMV infection. Memory B cell subsets, while exhibiting minimal age-related transcriptional reprogramming at homeostasis, demonstrate significantly altered responses to vaccine-induced perturbation in older adults. We further reveal that altered memory B cell responses correlate with memory CD4 T cell reprogramming towards a Th2-skewed state across age, providing insights into mechanisms of age-related immune dysregulation and unique targets for therapeutic intervention and disease prevention in older adults.

## Results

### Generation of a high-resolution healthy peripheral immune cell atlas for application across human immune studies

A critical step in understanding the immune landscape in healthy people across age is defining robust immune cell types within a tissue of interest. A number of references exist for labeling of human immune cells within peripheral blood using scRNA-seq datasets (Domínguez Conde et al. 2022; Terekhova et al. 2023; Hao et al. 2021), however none of these references collectively met our input criteria. These criteria included 1) use of many donors to account for immune composition heterogeneity, 2) large number of cells sequenced per donor to allow detection of rarer subsets, and 3) a broad age range of donors to account for age-specific variation. Thus, we began by building a new human immune cell atlas, based on scRNA-seq data we generated from peripheral blood mononuclear cells (PBMCs) of more than 100 healthy donors ranging from 11 to 65 years (yrs) of age (n=108, **Figure 1A**). Cohort details are provided in **Supplemental Table 1**. The cell numbers after quality control and doublet removal averaged more than 15,000 cells per donor (mean: 16,867 cells, range of 25-75 quartiles: 15074-18264 cells, sample-specific metrics are included in **Supplemental Table 1**), generating a final dataset of 1.82 million high-quality PBMCs from healthy people across age. From these data, we used a hierarchical label strategy, unsupervised clustering and distinct immune-based marker genes to define 9 cell subsets at level 1, 29 subsets at level 2 and 71 subsets at level 3, that further encompasses broad features of age, sex and cytomegalovirus (CMV) infection. (**Figure 1B, 1C**) Level 3 includes characterization of 35 T cell, 11 B cell, 7 monocyte, 6 natural killer (NK) and 12 other subsets including dendritic cells and hematopoietic precursors. (**Figure 1D**, see **Methods**). The number of cells used to generate this atlas allowed resolution of smaller, more unique subsets such as CD27^−^ TBX21^+^ effector B cells and the recently described population of circulating KLRC2^+^ CD8-alpha alpha T cells. (Thomson et al. 2023) Further details on our Human Immune Health Atlas and immune cell subsets can be found at https://apps.allenimmunology.org/aifi/resources/imm-health-atlas/. We utilized this high-quality, high-resolution human peripheral immune cell atlas for labeling cells in all subsequent scRNA-seq analyses presented here, providing consistent comparison of immune subsets across RNA and multi-modal single cell sequencing datasets.

**Figure 1.**
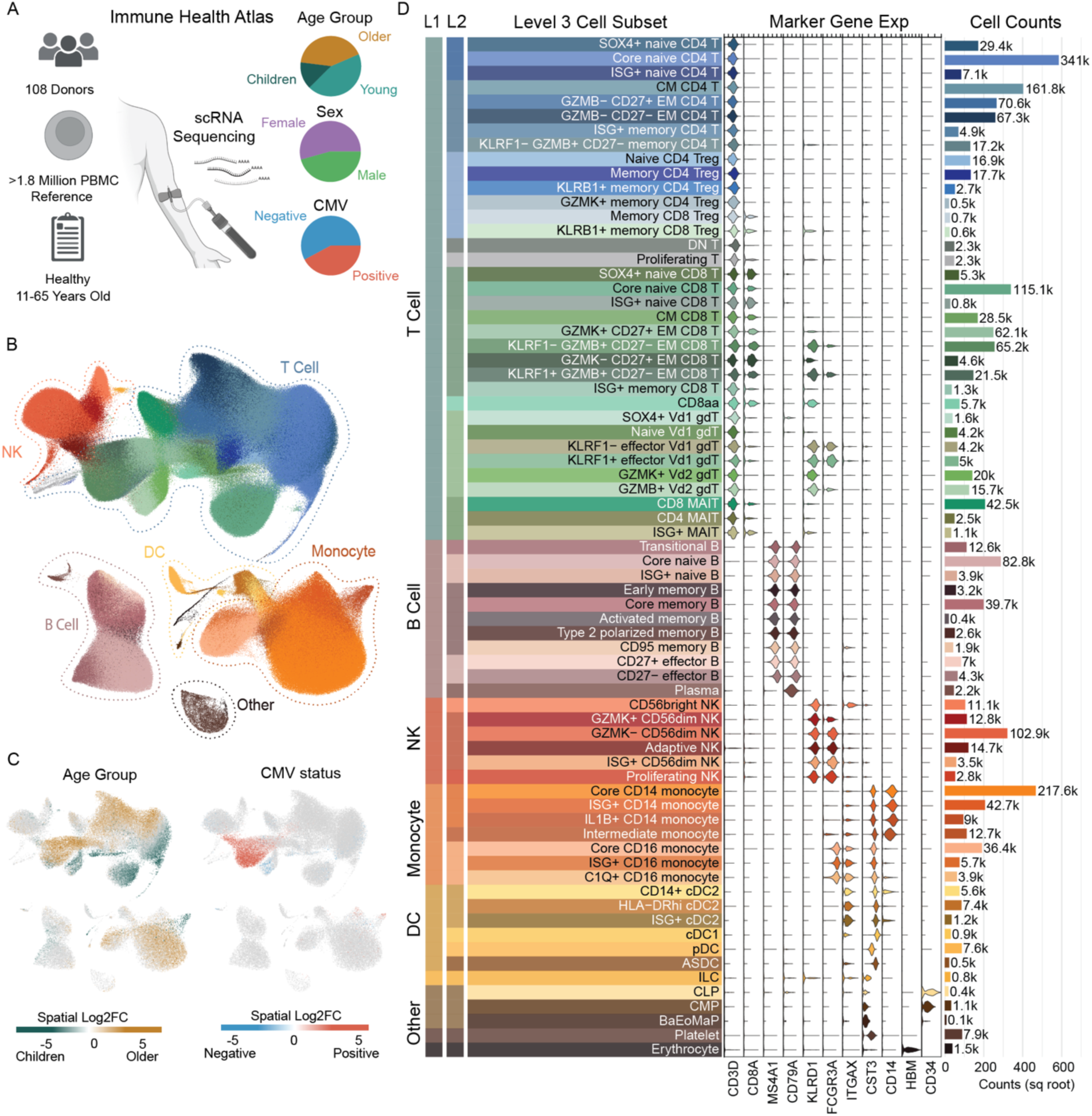
Generation of a high-resolution scRNA-seq atlas of peripheral immune cells from healthy children and adults. **A.** Overview of the Human Immune Health Atlas cohort (age range: 11-65 yrs; n=108) and final reference dataset. **B**. UMAP of immune cell subsets within the Atlas, highlighting major immune cell populations. **C**. Log_2_ fold change of clinical metadata features of age and CMV infection status compared using Milo differential abundance testing. Bronze is higher in older adults and Teal is higher in young adults. Red is higher in CMV^+^ people and blue is higher in CMV^−^ people. **D.** Marker gene expression and cell counts of the 71 immune cell subsets in level 3 of the Atlas. More details about this Atlas can be found at https://apps.allenimmunology.org/aifi/resources/imm-health-atlas/.

### Changes in the homeostatic programming of T cell subsets accumulate over the course of age, independent of systemic inflammation

To-date, there are limited longitudinal studies focused on understanding the dynamics of the immune cell landscape across healthy age, as these types of studies require two different time scales of “aging”; 1) comparison of donors of different ages and 2) repeat sampling of individual donors over time. To address this gap, we prospectively recruited a cohort of 49 young adults (25-35 yrs of age at enrollment) and 47 older adults (55-65 yrs of age at enrollment) and followed them longitudinally over the course of 2 years. (**Figure 2A**) During this time course, donors received 2 seasonal influenza vaccinations and up to 10 total blood draws. Vaccine-related blood draws were collected at 0, 7 and 90 days post-vaccination (“Flu Vax” series). A similar 0, 7 and 90 day time course was collected but with no vaccination administration (“No Vax” series), as well as additional “stand-alone” visits to account for seasonal variation. These time points were designed to enable comparison of age-related differences in the immune landscape that occur during homeostatic maintenance as well as during vaccine-induced perturbation. CMV serology was additionally performed to allow comparison of the impact of immune perturbation induced by chronic viral infection. PBMCs and plasma were collected at each blood draw for in-depth immune profiling assays that included scRNA-seq, spectral flow cytometry, O-link plasma proteomics and influenza-specific serology. Extensive clinical data was collected on donors at each blood draw and is available for detailed exploration at https://apps.allenimmunology.org/aifi/insights/dynamics-imm-health-age/vis/clinical/. Basic cohort demographics, including age, sex, CMV infection status, and sample information are detailed in **Supplemental Table 1**.

**Figure 2.**
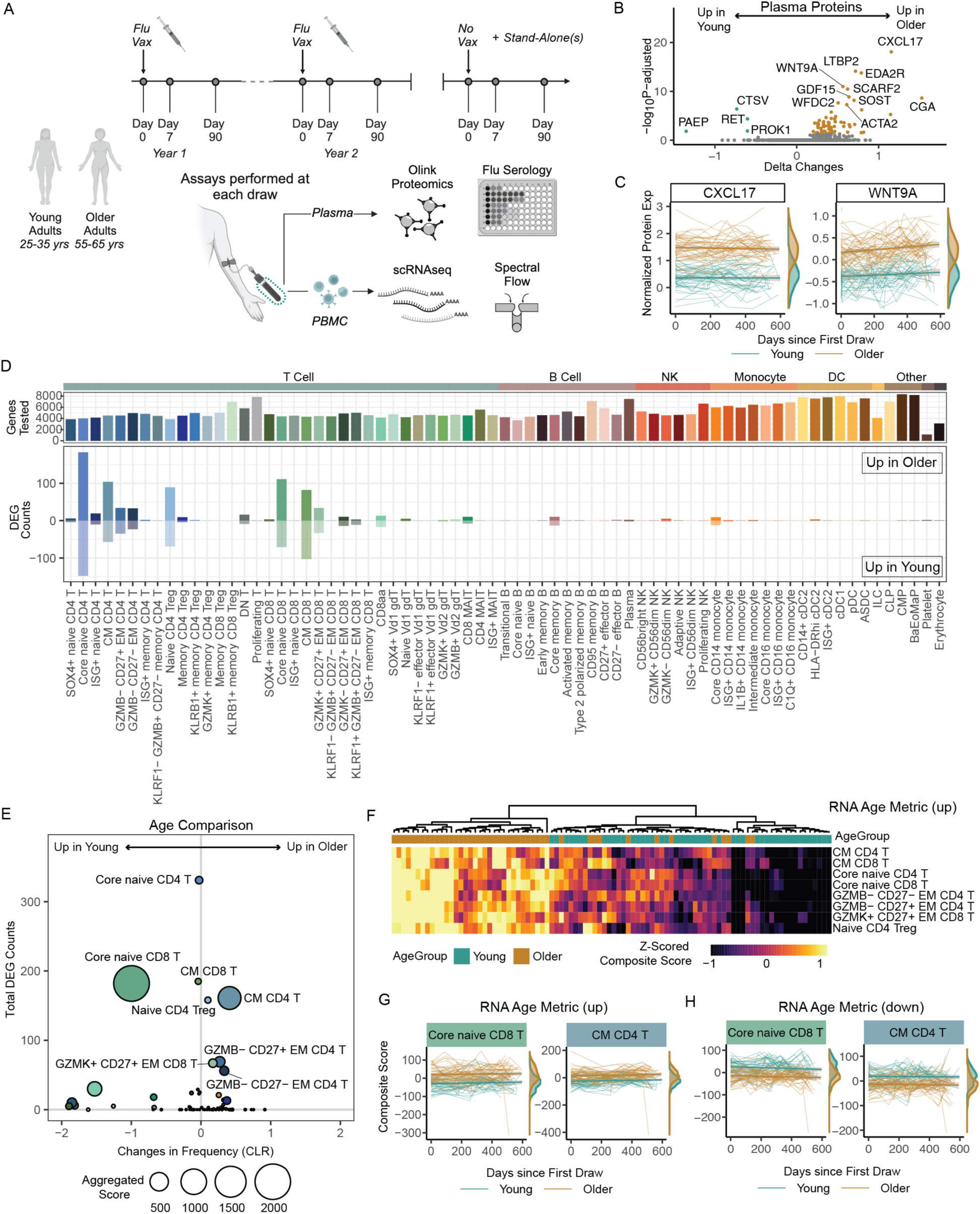
Maintenance of age-related alterations in the healthy human immune landscape over time. **A.** Overview of the longitudinal Sound Life cohort of healthy young (n=49) and older (n=47) adults. **B.** Volcano plot of the age-related protein expression differences in circulating plasma proteome at baseline (Flu Vax Year 1 Day 0). **C**. CXCL17 and WNT9A normalized protein expression (NPX) over time in young (teal) and older (bronze) adult plasma. Each donors’ samples are connected by a line. **D**. The number of differential expressed genes (DEGs) from DEseq2 analysis (log2fc >0.1 and p.adj<0.05) of immune cells subsets from young and older adults at ‘Flu Vax Year 1 Day 0’. **E.** Bubble plot comparison of the change in frequency (using centered log-ratio (CLR) transformation) and number of DEGs at ‘Flu Vax year 1 day 0’ between young and older adults. Bubble size shows a combined metric of change defined as −log10(p.adj from CLR freq comparison) x DEG_Counts. P.adj for CLR freq was determined using Wilcoxon rank-sum test with Benjamini–Hochberg correction. **F**. The RNA age metric, calculated as a composite score of the top upregulated DEGs for each subset with >20 DEGs, shown across each donor at Flu Vax Year 1 Day 0. **G.** RNA age metric (upregulated genes) in select subsets over time in young and older adults. Each donors’ samples are connected with a thin line. **H.** RNA age metric (downregulated genes) in select subsets over time in young and older adults. Each donors’ samples are connected with a thin line.

Age-related changes in the circulating proteome are well-described (Argentieri et al. 2023; Whiting et al. 2015), however there are conflicting results regarding the association between circulating markers of inflammation and age. Thus, we first investigated proteomic changes in our healthy adult cohort via our Olink dataset. We found 69 proteins differentially expressed at baseline between young and older adults (65 increased and 4 decreased with padj<0.05), including previously described markers including CXCL17 and WNT9A at baseline. (**Figure 2B, 2C**) (Argentieri et al. 2023) Notably, no significant increase in classic inflammatory proteins TNF, IL-6, IL-1B or a more recently described age-related marker IL-11 (Widjaja et al. 2024) were detected over time in older adults. (**Figure S1A**) The observed age-related alterations were also maintained over time, with a similar pattern of protein differences observed a year later. (**Figure S1B, S1C**) Thus, we find circulating hallmarks of healthy age persist in the absence of systemic inflammation.

We next interrogated the cellular landscape of healthy immune aging utilizing our scRNA-seq dataset. To build the reference dataset for this study, all scRNA-seq data collected from our cohort was compiled and cells were labeled via our new Human Immune Health Atlas. After cell label transfer, post-transfer doublet exclusion, and clean-up, the resulting longitudinal immune health dataset consisted of >13.7 million PBMCs. Similar to the atlas dataset, the cell numbers per sample averaged more than 15,000 cells per donor per time point (mean: 15,886 cells, range of 25-75 quartiles: 14,153-18,032 cells, sample-specific metrics are included in **Supplemental Table 1**). The final dataset includes more than 3 million T cells, 1.2 million B cells, 1.1 million NK cells, 2.4 million monocytes, 123,020 dendritic cells, and 10,431 hematopoietic precursors, building a rich resource to interrogate age-related immune changes at high-resolution in many immune cell subsets simultaneously.

A consistent hallmark of immune aging is the loss of naïve CD8 T cells. We began by examining cellular composition as a key feature of the aging process in this reference dataset. We found significant age-related compositional changes in 16 immune cell subsets, including decreased frequencies of core naïve CD8 T cells at baseline (padj = 8.6e-12) in older adults that is a hallmark feature of immune aging. (**Supplemental Table 2**) To further explore features of homeostatic immune aging, we next focused on examining transcriptional profiles across all 71 immune cell subsets defined by our Human Immune Health Atlas. For these analyses, we performed pseudo-bulk differential gene analyses comparing young and older adults at baseline (“Flu Year 1 Day 0”), controlling for sex and CMV as potential confounding factors. We found that T cell subsets were the main immune cells that exhibited transcriptional changes with age. (**Figure 2D**) Consistent with previous T cell-focused studies (Thomson et al. 2023; Moskowitz et al. 2017), T cell subsets early in differentiation showed the highest number of age-related differentially expressed genes (DEGs), with naive > central memory > effector memory subsets. Of note, few transcriptional changes were observed in other, non-T cell subsets and many subsets exhibited no age-related changes in their transcriptome. To confirm this observation, we ran a cross-sectional analysis on samples collected one year later (“Flu Year 2 Day 0”) and found similar results. (**Figure S1D, S1E**) Age-related changes were also distinct from those found in immune subsets when directly assessing broad immune perturbations caused by CMV infection and influenza vaccination. (**Figure S1F**) Indeed, many transcriptional differences in B cell subsets were found with vaccination, highlighting that the lack of age-related differences in this adaptive immune compartment is unlikely a technical artifact due to lower B cell numbers in the peripheral blood. While T cell subsets also demonstrated the most significant changes in frequencies with age, the transcriptional changes did not directly correspond with compositional changes in the subsets. (**Figure 2E**) Indeed, core naive CD4 T cells show the most transcriptional changes (N=331 DEGs) with no significant difference in frequency (padj=0.65), whereas core naive CD8 T cells show both transcriptional (N=182 DEGs) and frequency changes with age. Thus, during homeostasis, T cells are the peripheral immune cell subset most transcriptionally and compositionally altered by age, with few transcriptional changes observed in other innate and adaptive immune compartments.

The homeostatic aging process has been linked with transcriptional variation however the actual stability of immune cell programming across age is unknown. To further determine the stability of age-related transcriptional changes in T cells, we analyzed the longitudinal transcriptional changes (over the course of 600 days) in the 8 immune cell subsets with the most differentially expressed genes. These subsets included core naive CD8 T cells, core naive CD4 T cells, central memory CD8 T cells (N=185 DEGs), GZMK+CD27+ effector memory CD8 T cells (N=67 DEGs), naive CD4 Tregs (N=158 DEGs), central memory CD4 T cells (N=161 DEGs), GZMB-CD27-effector memory CD4 T cells (N=56 DEGs) and GZMB-CD27+ effector memory CD4 T cells (N=69 DEGs). To interrogate the overall maintenance of transcriptional profiles with age, we developed upregulated and downregulated RNA-based composite scores as summary metrics of age-related differential gene expression specific to each of these subsets. (**Figure S2A**) Applying these metrics to each donor, we found consistent age-related transcriptome changes in young and older adults (adjusted p<0.05 for all subsets). (**Figure S2B**) There was also a significant correlation between age metric across each of the 8 T cell subsets within an individual (**Figure 2F**), implying that transcriptional changes occur uniformly across the T cell compartment of an individual with age. Moreover, older adults consistently maintained differential age metrics compared with young adults over a 2-year period. (**Figure 2G, 2H**, **Figure S2C, S2D**). These data collectively indicate that age-related transcriptional differences are maintained over time in age-susceptible T cell subsets.

To further collectively confirm that these proteomic, compositional and transcriptional changes accumulate over the course of homeostatic aging, we acquired a second cross-sectional cohort of paired PBMC and serum samples from healthy adults (n=234), with a continuous age range from 40 to 90+ years of age (Whiting et al. 2015), and again performed deep immune profiling. (**Figure 3A**) Basic cohort demographics, including age, sex, and CMV infection, as well as sample assay information are detailed in **Supplemental Table 1**. Proteomic analysis revealed circulating proteins altered with healthy age continuously accumulate across the aging spectrum in the absence of systemic inflammation, evidenced by the lack of association with TNF, IL6, IL1B and IL11 with chronological age. (**Figure 3B**) We further utilized the scRNA-seq data to build a follow-up reference of more than 3.2 million PBMCs of similar quality as our original datasets (**Figure 3C**; see Methods). Initial examination of the 71 cell subsets confirmed the expected hallmarks of immune aging, including a decrease in core naive CD8 T cells (slope: −7.6, pval=5.3e-26) and a modest, although not significant, increase of CM CD4 T cells (slope: 2.12, pval=1.4e-01) across age. (**Figure 3D, 3E**) To better understand the dynamics of the transcriptional programming across age, we applied our previously described summary metrics of age-related differential gene expression (‘transcriptional age” metric). We found a continuous increase in the age metric for upregulated genes across the 8 age-susceptible subsets from people 40 to 65 years of age. (**Figure 3F**) Notably, the upregulated RNA age metric consistently plateaued after 65 years of age across all subsets. Conversely, the age metric for down-regulated genes demonstrated the opposing pattern, remaining relatively stable from 40-65 then rapidly decreasing from 65-90 yrs of age. (**Figure 3G**) This suggests that up- and down-regulated genes within immune cells have different dynamics in adults above 65 years of age and may be distinct from those occurring between 40-65 years of age. To further investigate the functional implications of these changes, we assessed the overlap between DEGs in these 8 subsets to find commonly targeted genes; identifying 26 gene down-regulated and 36 gene up-regulated with age that were consistent in at least 4 subsets. Common age-related genes included increasing genes associated with effector differentiation (e.g., *PTGER2*) and intracellular signaling responses (e.g., *SESN3*, *ANHAK*) (**Figure 3H**), as well as decreasing genes associated with cell state polarization (e.g., *IL16*) and apoptosis (e.g., *STK17A*). (**Figure 3I**) Collectively, these data demonstrate that age-related changes in the immune landscape affect T cell subsets early in differentiation and coordinated changes occur progressively over the course of age, independent from markers of systemic inflammation.

**Figure 3.**
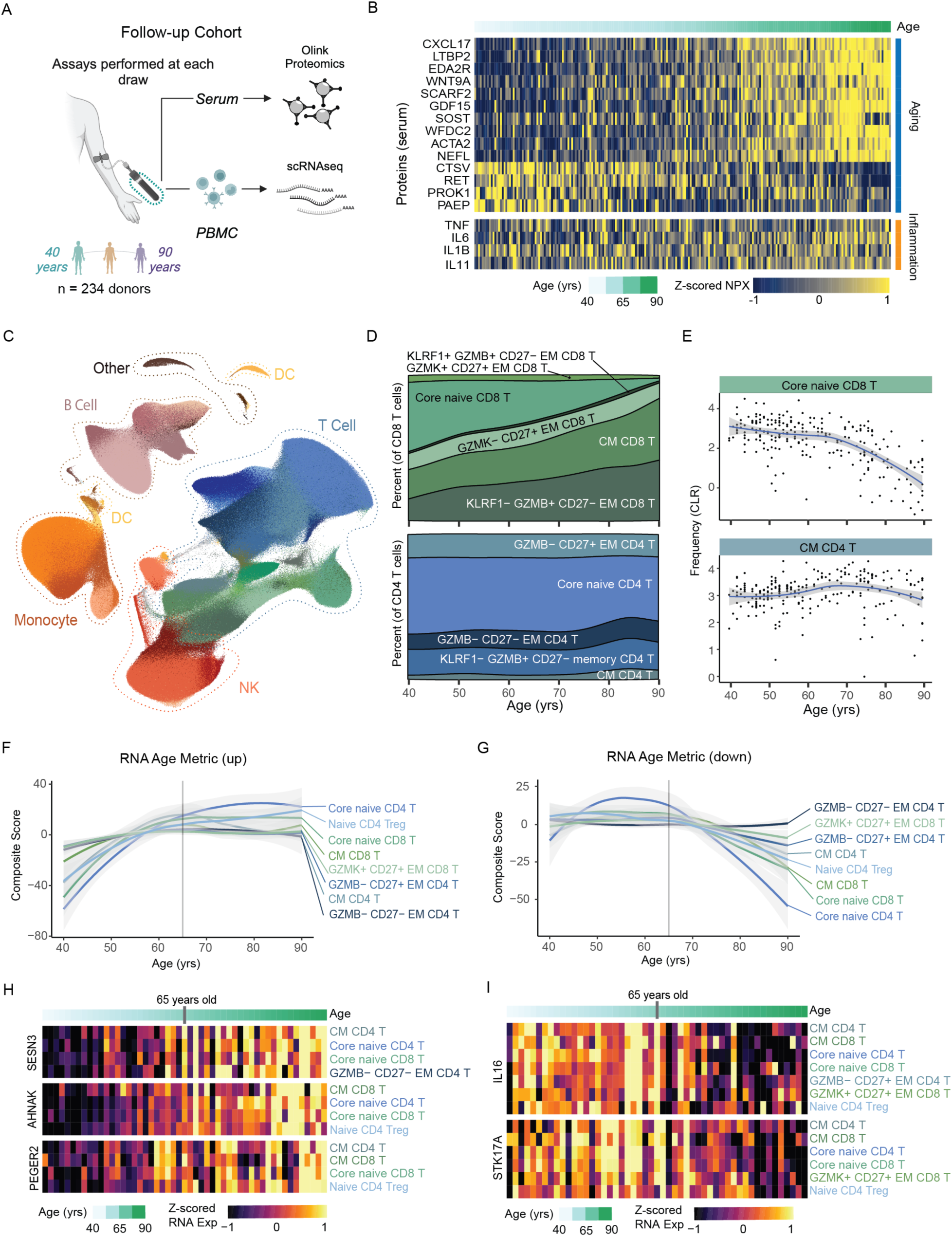
Dynamics of the healthy human immune landscape across age. **A.** Overview of our follow-up cohort of healthy adults (n=234) ranging from 40 - 90 years of age. **B.** Normalized protein expression (NPX) of select age- and inflammation-related serum proteins in our follow-up cohort, with donors ordered by age. **C.** UMAP of scRNAseq data generated from our follow-up cohort, totaling a final reference dataset of 3.2 million PBMCs. **C.** Distribution of immune cells by sex and CMV infection status with the UMAP. **D.** Composition of CD8 and CD4 T cell compartment across age. **E.** Frequencies (using centered log-ratio (CLR) transformation) of select T cell subsets within PBMCs across age. Regression line shown with 95% confidence intervals in gray. **F.** The average RNA Age Metric (upregulated genes) for the top age-impacted immune cell subset, shown across age. Regression line shown with 95% confidence intervals in gray. **G.** The average RNA Age Metric (downregulated genes) for the top age-impacted immune cell subset, shown across age. Regression line shown with 95% confidence intervals in gray. **H-I.** Heatmap of mean RNA expression by age in the follow-up cohort for select **H.** up-regulated and **I.** down-regulated genes identified from initial DEG analysis.

### The immune landscape induced by chronic CMV infection is stable over time and distinct from aging

Cytomegalovirus (CMV) is thought to contribute to low-grade inflammation by inducing chronic immune stimulation over time. However, the impact of age on the cellular and molecular landscape induced by CMV infection has yet to be fully elucidated. To assess the overall impact of CMV on the immune landscape of our healthy adult cohort, the cell frequencies and transcriptomes of the 71 immune cell subsets were compared in CMV^+^ and CMV^−^ individuals (CMV^+^ n = 44; CMV^−^ n = 52). We found 6 cell subsets to be significantly impacted by CMV infection (i.e., with greater than 5 DEGs and cell frequency differences of adj p<0.05). (**Figure 4A**) In particular, we found increased frequencies of KLRF1^+/-^GZMB^+^ EM CD8 T cells (KLRF1^+^ padj=0.002; KLRF1^−^ padj =3.9e-10), KLRF1^−^GZMB^+^ EM CD4 T cells (padj=1.5e-18), GZMK^+^ CD27^+^ EM CD8 T cells (adjusted p= 0.02), KLRF1^+^ γδ T cells (padj = 1.8e-07) and adaptive NK cells (padj = 1.2e-08) in CMV^+^ compared with CMV^−^ individuals. Immune cell subsets with CMV-related cell frequency increases also displayed more prominent transcriptional changes, including KLRF1^+/-^GZMB^+^ EM CD8 T cells (KLRF1^+^ N=23 DEGs; KLRF1^−^ N=56 DEGs), KLRF1^−^GZMB^+^ EM CD4 T cells (N=8 DEGs), and adaptive NK cells (N=63 DEGs). These findings were confirmed in our larger, follow-up cohort (CMV^+^ n=136; CMV^−^ n=98). (**Figure S3A**) Thus, we determined specific innate and adaptive immune cell subsets commonly expanded in chronic CMV infection, regardless of age.

**Figure 4.**
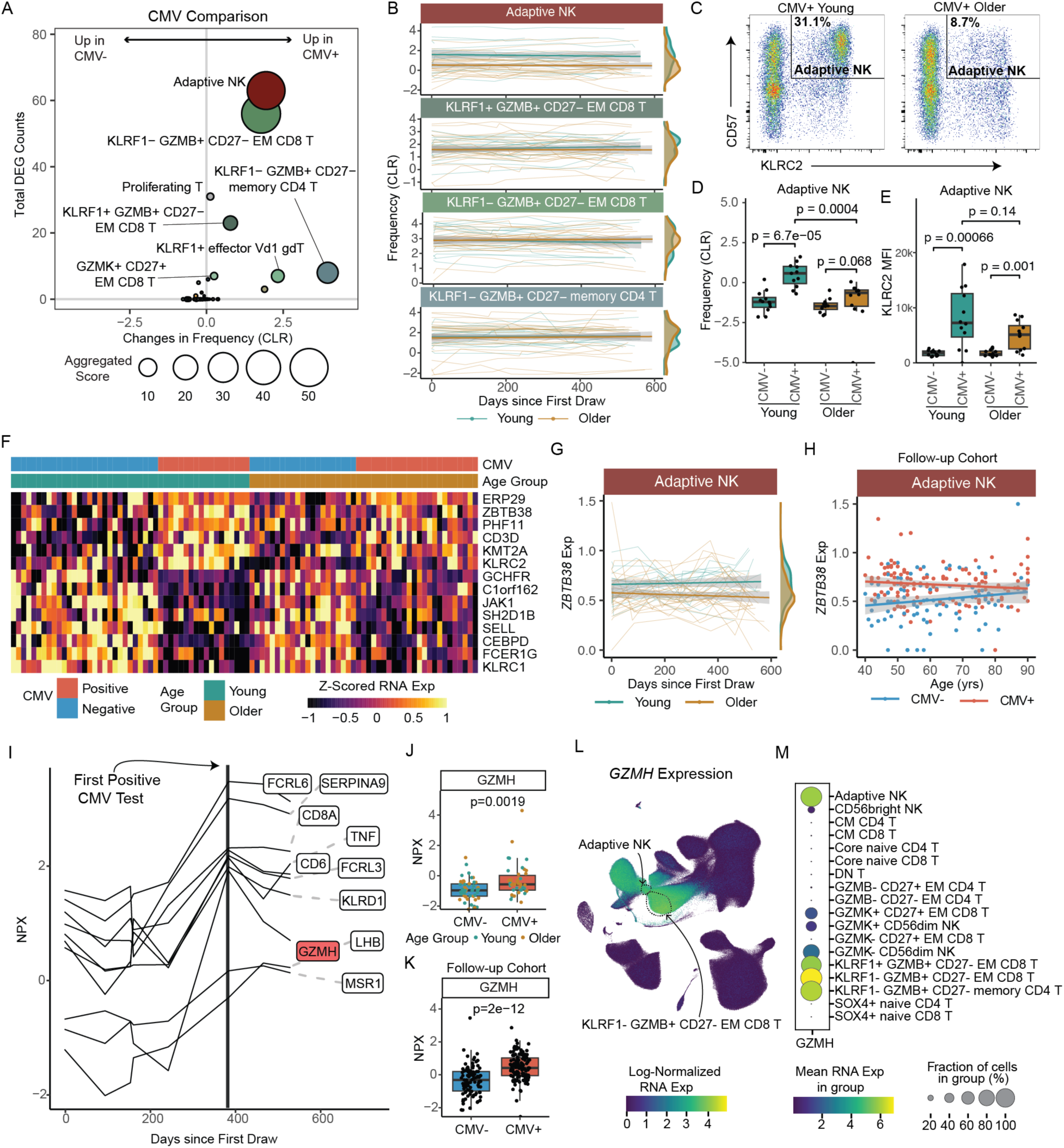
Distinct impact of CMV infection and age on the immune landscape. **A.** Bubble plot comparison of the change in frequency (using centered log-ratio (CLR) transformation) and number of DEGs at ‘Flu Vax year 1 day 0’ between CMV^+^ (n=44) and CMV^−^ (n=52) adults. Bubble size shows a combined metric of change defined as −log10(p.adj from CLR freq comparison) x DEG_Counts**. B.** Select subset frequencies in PBMCs shown over time. Teal dots are young adults. Bronze dots are older adults. Regression line shown. **C.** Representative flow plots of CD57 and NKG2C (*KLRC2* gene) expression within NK cells. Adaptive NKs cells are defined as CD57^+^NKG2C^+^ NK cells. **D.** Adaptive NK cell frequencies and **E.** NKG2C (KLRC2) MFI expression on adaptive NKs comparing young CMV^−^ (n=12), young CMV^+^ (n=12), older CMV^−^ (n=12) and older CMV^+^ (n=12) adults from spectral flow cytometry analysis. P-values calculated using unpaired Wilcoxon test. **F.** Heatmap of mean RNA expression levels from upregulated DEGs in adaptive NKs of CMV^+^ young adults across all individuals in longitudinal cohort (n=96). **G**. *ZBTB38* expression in adaptive NKs (left panel) CMV^+^ young (n=18) and older (n=24) adults, shown over time (up to 600 days after first blood draw). Teal dots are CMV+ young adults. Bronze dots are CMV^+^ older adults. Regression line shown. **H.** *ZBTB38* expression in adaptive NKs (right panel) CMV^+^ (red, n=136) and CMV^−^ (blue, n=98) adults across age in our follow-up cohort. **I.** Normalized expression (NPX-bridged) of plasma proteins in one young adult who converted from CMV^−^ to CMV^+^ over the course of our study. Proteins were considered significant if they had a 1.5 or greater fold change pre- to post-conversion. **J**. Normalized expression of GZMH protein in plasma of CMV^−^ and CMV^+^ individuals from our longitudinal cohort. Young and older adults are delineated by circle and squares, respectively. **K.** Normalized expression of GZMH protein in serum of CMV^−^ and CMV^+^ individuals from our follow-up cohort. P-value for J and K were determined by Wilcoxon rank-sum test. **L.** UMAP of GZMH RNA expression in PBMCs from all individuals in our longitudinal cohort. **M**. Dot plot of GZMH RNA expression in NK, CD4 T cell and CD8 T cell subsets.

We next compared the age-associated stability of compositional changes observed in CMV^+^ people, focused on these 6 commonly expanded immune cell subsets. No age-related differences in the frequencies of the more terminal-like effector T cell subsets, KLRF1^+/-^GZMB^+^ EM CD8 T cells, in CMV^+^ adults were observed both at baseline and over a two-year period (**Figure 4B**, **Figure S3B**), possibly due to the relatively ‘younger’ nature of our healthy cohort (less than 65 yrs of age). There was, however, a significant decrease in adaptive NK cells (padj=0.01) in CMV^+^ older adults compared with CMV^+^ young adults, which was maintained over a two-year period (**Figure 4B**). To confirm this age-related decrease of adaptive NK cells, we performed spectral flow cytometry on 24 young (CMV^+^ n=12; CMV^−^ n=12) and 24 older (CMV^+^ n=12; CMV^−^ n=12) adults. While both CMV^+^ young and older adults showed expanded adaptive NKs compared with CMV^−^ individuals, adaptive NK cells (defined as KLRC2^+^CD57^+^CD56dimCD16^+^ cells (Dogra et al. 2020; Lopez-Vergès et al. 2011)) had a lower frequency in older CMV^+^ adults compared to young CMV^+^ adults. (**Figure 4C, 4D**) We also noted a trending decrease in expression of KLRC2 (a.k.a., NKG2C) on adaptive NK cells from CMV^+^ older individuals. (**Figure 4E**) Thus, age modestly impacts the composition of the CMV-mediated immune landscape in healthy adults.

We then evaluated the impact of CMV infection on age-related differences in the transcriptome of age-susceptible T cell subsets (identified in **Figure 2**), using our RNA age metric. Consistent with CMV infection mainly impacting more terminal effector T cell subsets, no significant differences in the age-related transcriptional profiles of these 8 T cell subsets between CMV^+^ and CMV^−^ young or older adults were found (**Figure S3C**, p>0.05 for all 8 subsets), indicating that chronic viral infection does not accelerate transcriptional age in T cell subsets early in differentiation. We assessed how age impacts the transcriptional changes induced by chronic CMV infection in all immune cell subsets, comparing CMV-induced gene expression in young and older adults separately. From these analyses, we found that adaptive NKs and KLRF1^−^GZMB^+^ EM CD8 T cells displayed the most transcriptional changes with CMV infection in young adults, however these changes were much less pronounced in older adults at baseline. (**Figure S3D**) The number of CMV-induced DEGs in adaptive NKs were also found to be consistently reduced in older adults one year later (Flu Year 2 Day 0). (**Figure S3E**) Moreover, a subset of CMV-induced genes in adaptive NKs of young adults exhibited overall lower expression in older adults, including *KLRC2*. (**Figure 4F**) In particular, *ZBTB38*, a newly defined marker gene of the adaptive NK population (Rebuffet et al. 2024), was decreased, both consistently over time in CMV^+^ older adults compared with CMV^+^ young adults (**Figure 4G**), as well as decreasing in expression in CMV^+^ subjects across age in our follow-up cohort, reaching similar expression to CMV^−^ subjects in advanced age. (**Figure 4H**) Thus, the changes in the transcriptional landscape induced by CMV infection are relatively independent of age, however the adaptive NK subset uniquely exhibits a stably altered transcriptional state suggestive of lower activation in older adults.

CMV infection may have systemic effects on the immune system as well, thus we evaluated the impact of CMV and age on the circulating proteome. Although we did not find any significant different proteins between CMV^+^ and CMV^−^ individuals at a global level, we took advantage of the one young adult that seroconverted from CMV^−^ to CMV^+^ during our longitudinal study to identify select proteins that are induced by CMV infection. CMV-induced proteins were determined via strict criteria of being expressed at low levels during the CMV uninfected phase but had a persistent increase after infection of at least 1.5-fold higher than the highest pre-infection level. We identified 10 proteins that met these criteria. (**Figure 4I**) We next evaluated whether there was a significant difference in expression levels of these 10 proteins in CMV^+^ vs. CMV^−^ individuals from our full longitudinal cohort. We found that 4 of the proteins (FCRL6, FCRL3, KLRD1, GZMH) were significantly increased in chronic CMV infection (**Figure 4J**, *not shown)*, independent of age. Additionally, higher expression of GZMH in CMV^+^ individuals was maintained over time and showed significant CMV-related differences in our follow-up cohort, again independent of age. (**Figure 4K**) To further link these proteomic changes back to the observed changes in cellular composition, we evaluated which immune cell subsets transcriptionally express GZMH. GZMH was expressed the highest in GZMB^+^ T cell subsets and adaptive NKs (**Figure 4L, 4M**), linking CMV-related immune cell expansion with the circulating proteome. Together, these data demonstrate that CMV infection stably alters the immune cell landscape in adults, with age primarily altering adaptive NK cells in healthy CMV^+^ adults.

### Age-related alterations in B cell responses to vaccine-induced perturbation are maintained over time

It is well-known that age also has a major impact on the ability of an individual to make effective, long-lasting antibody responses to vaccination, linked to alterations in the B cell compartment (Wang et al. 2019; Frasca et al. 2016), however less is known about the impact of age on broader B cell responses to vaccine-induced perturbation (i.e., bystander activation) in healthy adults. To study these broad responses, we first binned individuals into their specific “flu vaccine year” to account for variation in seasonal flu vaccine composition (e.g., flu strains, adjuvants) that may impact cell responses, identifying 26 young and 42 older adults receiving the same vaccine in the year. (**Figure 5A**) Then we assessed whether our sub-cohort was generally representative of other aging cohorts, examining total level of influenza vaccine-specific IgG antibodies as well as the functional capacity of vaccine-specific antibodies using a custom hemagglutination inhibition (HAI) MSD assay at day 0 and day 7 post-vaccination. We additionally evaluated antibody responses against a vaccine neo-antigen (B/Washington strain; **Figure 5B, 5C**) and a vaccine recall antigen (B/Phuket strain; **Figure 5D, 5E**). Consistent with previous aging studies, the functional antibody response (HAI) to a recall antigen was significantly lower in older adults. (**Figure 5E**) However, the neo-antigen HAI responses were similar between young and older adults (**Figure 5C**), as were total vaccine-specific IgG levels for both neo- and recall antigens. (**Figure 5B, 5D**) Thus, these data consistently demonstrate that age impacts vaccine-specific antibody responses and uniquely indicate that memory, but not naïve, B cell responses are preferentially impaired with age.

**Figure 5.**
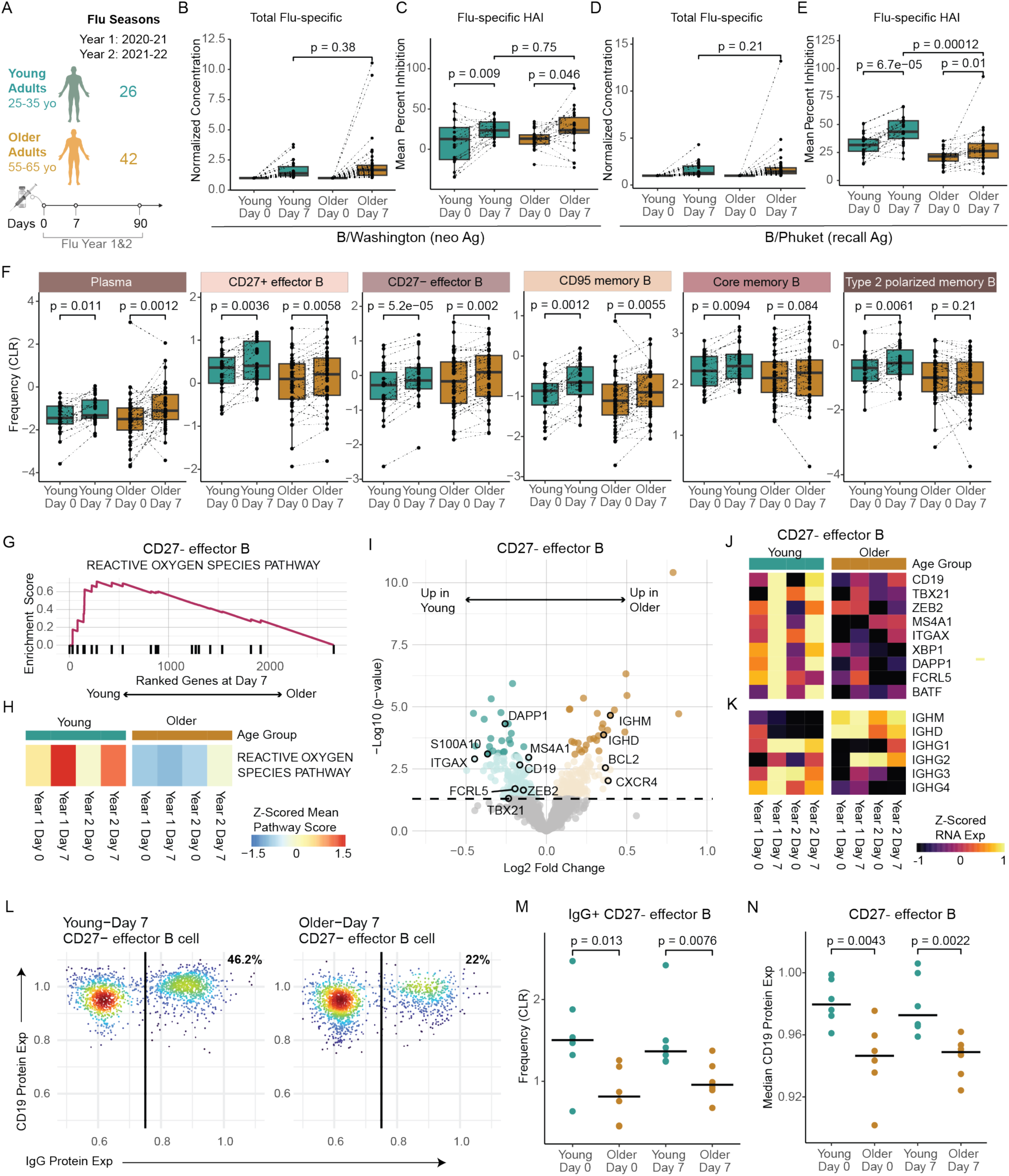
Age-associated B cell responses to the influenza vaccine. **A.** Number of samples and sampling timepoints across 2 flu seasons in the young and older adult cohorts that received the same seasonal vaccines. **B-C.** B/Washington and **D-E.** B/Phuket flu-specific total IgG antibody expression (**B, D**) in plasma compared to expression at baseline (Day 0) for both young (n=26) and older (n=42) adult cohorts in Flu Vax Year 1 and Mean percent inhibition of flu hemagglutinin (HA) antigen as determined by the HAI assay (**C, E**) for both young (n=21) and older (n=22) adult cohorts at days 0 and 7 in Flu Vax Year 1. P-values were calculated using Wilcoxon’s signed-rank test (paired) for the comparison between Day 0 and Day 7, and using the Wilcoxon rank-sum test for all other comparisons. **F.** Peripheral memory and antibody-secreting B cell population frequency changes in young (teal) and older adult (bronze) cohorts pre-vaccination (Day 0), and post-vaccination (Day 7). P-vals determined by Wilcoxon’s signed-rank test (paired). **G.** Enrichment plot for the top Hallmark pathway in CD27-effector B cells when comparing Day 7 transcriptome between young and older adults after gene set enrichment analysis. **H.** Sample level enrichment analysis scores for the Hallmark Reactive Oxygen Species Pathway at each timepoint for CD27-effector B cells in young and older adults. **I.** Volcano plot of DEGs for CD27-Effector B cells between the age cohorts at Day 7 post-vaccination. Highlighted genes are those previously shown to define a flu-specific effector B cell subset in a vaccinated adult cohort. Dark teal and bronze dots signify significantly different genes, and light-colored dots indicate nominal significance, while gray dots indicate no significance between age cohorts. **J-K.** Longitudinal expression of selected genes by CD27-effector B cell subset, averaged for each age cohort at each timepoint. **L**. Representative flow cytometry plot of CD19 and IgG protein expression on CD27-effector B cells in young and older adults at day 7 post-vaccination, based on flow cytometry analysis of 6 young and 6 older adult subjects that overlap with the scRNA data cohort. **M.** CLR-transformed frequency comparison of surface IgG+ CD27-effector B cells pre- and post-vaccination in young and older adults, as determined by flow cytometry. P-values were determined by Wilcoxon rank sum test with the alternative hypothesis ‘less’. **N.** Median surface CD19 protein expression comparison on CD27-effector B cells pre- and post-vaccination in young and older adults, as determined by flow cytometry. P-values were determined by Wilcoxon rank sum test with the alternative hypothesis ‘less’.

In our previous analyses, we observed memory B cell subsets had the most transcriptional changes after vaccination. (**Figure S1F**) Thus, we utilize our high-resolution scRNA-seq dataset to further explore the impact of vaccine-induced perturbation on the composition and transcriptome of these subsets with age. We assessed age-related frequencies of plasma cells pre- and post-vaccination, as a classic cellular read-out of vaccine-specific responses in the periphery. Like the total flu-specific IgG responses, we found both young and older adults exhibited similar increased in plasma cells post-vaccination. (**Figure 5F**) We also observed similar increases in frequencies for multiple memory B cell subsets, including CD27^−^ effector B cells, CD27^+^ effector B cells and CD95^+^ memory B cells, independent of age. However, core memory and type 2 polarized B cell subsets had reduced expansion in older adults. (**Figure 5F**) We further confirmed these frequency changes via flow cytometry analysis on a subset of donors, demonstrating strong correlations between RNA- and flow-based B cell subset frequencies. (**Figure S4A**) Thus, while there was no alteration in the vaccine-specific expansion of plasma cells with age, we observe a broader, age-related limitation in expansion of memory B cell subsets with age.

To further assess broad memory B cell responses to vaccination, we interrogated transcriptional profiles at day 7 post-vaccination for enrichment of activation-related signaling pathways. Plasma cells exhibited pathway enrichment indicative of higher activation, whereas core memory B cells demonstrated lower MYC-related activation in older adults that aligned with their limited expansion with age. (**Figure S4B**) Similar to core memory, CD27+ effector B cells also lower enrichment in pathways associated with activation in older adults (e.g., MYC pathway, adjusted p value: 3.68-08, NES: 2.02). (**Figure S4C, S4D**) CD27-effector B cells also exhibited reduced ROS-associated signaling in older adults (adjusted p value: 0.04, NES: 1.77; **Figure 5G**), which has previously been linked to flu vaccine responsiveness. (Nellore et al. 2023) This reduction was observed at multiple different time points over the course of vaccination. (**Figure 5H**) Moreover, this effector B cell subset in older adults shows lower expression of genes associated with functional effector memory B cells (Nellore et al. 2023) (**Figure 5I, 5J**), including lineage-defining genes *FCRL5*, *CD19*, *MAS4A1*, and *ITGAX*, as well as activation genes like *ZEB2*, *TBX21*, *XBP1*, *S100A10*, *DAPP1*, and *BATF*. Collectively, these data indicate that older adults have broad alterations in the compositional and transcriptional B cell responses to vaccine-induced perturbation.

A key feature of effective B cell responses to flu vaccination is the production of class-switched immunoglobulin G (IgG) antibodies. Thus, we assessed whether the CD27-effector B cell subset displayed altered isotype composition post-vaccination. Notably, older adults displayed significantly lower expression of IgG genes (e.g., *IGHG1)* as well as higher *IGHD* and *IGHM* expression than young adults in CD27-effector B cells. (**Figure 5K**) The reduction in *IGHG* genes was found in both year 1 and year 2 flu vaccination series, demonstrating that there is a consistent decline in class-switch responses overall with age. To further confirm these data, we assessed IgG expression by the effector memory B cell subset via spectral flow cytometry using the B cell annotation strategy from Glass, et. al., 2020. (Glass et al. 2020) We found that frequency of IgG+ CD27-effector B cells was significantly lower in older adults both at 0 and 7 days post-vaccination than young adults (**Figure 5L, 5M**). Moreover, the CD27-effector population also expressed less surface CD19 in older adults (**Figure 5N**), corresponding to their age-related gene expression profiles. Thus, CD27-effector B cells, an important subset in response to flu vaccination, display lower IgG class-switching and express less lineage-defining surface markers with age. Together, these data reveal that multiple memory B cell subsets have altered age-related responses to vaccine-induced perturbation that could play a role in reduced antibody functionality in older adults.

### Accumulation of a Th2-like state in memory CD4 T cells is associated with age-related B cell dysregulation

Poor memory B cell activation, altered immunoglobulin class switching and reduced vaccine-specific antibody functionality in older adults suggests there may be alterations in memory T cell helper capacity with age. However, our initial flow cytometry-based analysis found no differences in the frequencies of activated T follicular helper cells (defined as ICOS^+^CD38^+^PD1^+^CXCR5^+^ CD4 T cells) at day 7 post-vaccination between young and older adults. (**Figure S5A**) These data, in tandem with similar frequencies of vaccine-induced plasma cells, indicate that the general process of antigen-specific activation and expansion are likely maintained with age. Thus, we determine if age-related molecular reprogramming of CM CD4 T cells, which are one of the most transcriptionally altered memory T cell subsets with age (**Figure 2**), could influence the type of helper functions these cells provide to B cells. Initial analyses of the overall ‘helper’ programming found that the Tfh transcriptional signature within the CM CD4 T cells decreased with age, (**Figure S5B**) with a main Tfh-defining marker CXCR5 expression decreased in CM CD4 T cells from older adults. (**Figure S5C**) This decrease was confirmed by a consistent loss of CXCR5 expression across age in our follow-up cohort. (**Figure S5D**) We next used CellPhoneDB (Efremova et al. 2020) to interrogate receptor-ligand interactions. (**Figure 6A**) Notably, we found that CM CD4 T cells from older adults displayed reduced predicted interactions with core memory B cells, including a key complex for B cell activation CD40LG:CD40. (**Figure 6B**) This reduced interaction exhibited similar decreases in these receptor-ligand interactions at year 2 (**Figure S5E**) and stemmed from a loss of CD40LG in CM CD4 T cells (**Figure S5F**), but not CD40 in core memory B cells, in older adults. (**Figure S5G**) This was confirmed in our follow-up cohort, which showed steadily decreasing expression of CD40LG in CM CD4 T cells, but not CD40 in core memory B cells, across age. (**Figure 6C**) Thus, CM CD4 T cells exhibit a lower transcriptional propensity for providing direct help to memory B cells with age.

**Figure 6.**
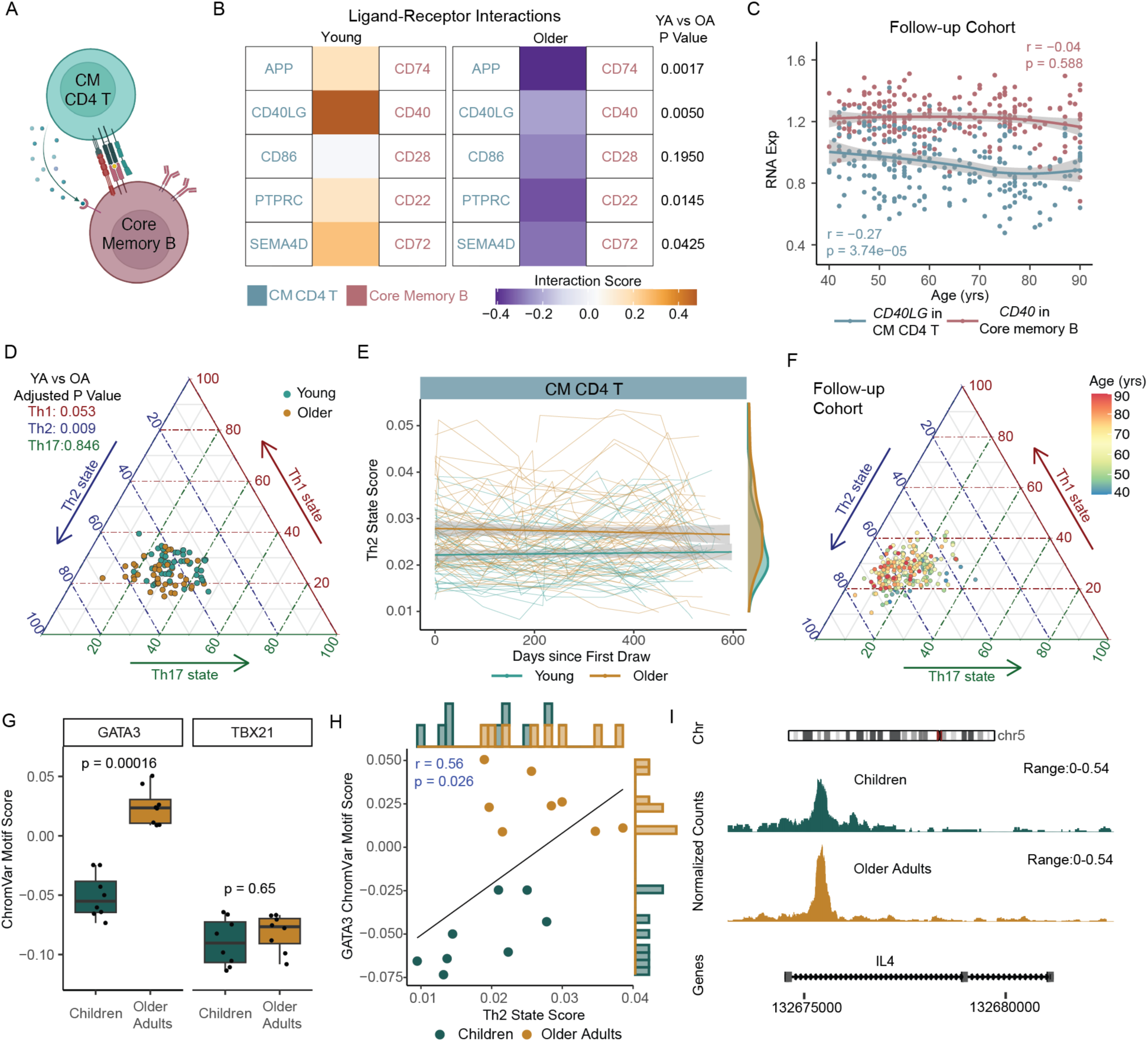
Accumulation of an altered transcriptional state in central memory T cells with age. **A.** Graphical representation of T cell and B cell interactions. **B.** Receptor-ligand interaction prediction between CM CD4 T cells and core memory B cells in young (n=47) and older (n=49) adults from a single time point (Flu Year 1 Day 0). **C.** CD40LG in CM CD4 T cells and CD40 in core memory B cells across age in our follow-up cohort (n=234). Regression line shown with 95% confidence intervals in gray. **D.** Triangle plots of Th1-, Th2- and Th17-cell state scores in CM CD4 T cells from young (teal) and older (bronze) adults. **E.** Th2 cell state scores in CM CD4 T cells over time in young (teal) and older (bronze) adults. Regression line shown with 95% confidence intervals in gray. **F.** Triangle plots of Th1-, Th2- and Th17-cell state scores in CM CD4 T cells from our follow-up cohort (n=234). **G.** GATA3 and TBX21 transcription factor (TF) activity based on Chromvar analysis of TEA-seq data in CM CD4 T cells from children (n=8) and older adults (n=8). **H**. Spearman correlation between GATA3 TF activity and Th2 cell state score in CM CD4 T cells from the TEA-seq dataset (n=16). **I.** Chromatin accessibility tracks of the *IL4* gene region in CM CD4 T cell subsets, showing normalized read coverage.

In addition to direct interactions, T cells can mediate B cell responses through indirect, cytokine-mediated interactions based on their help state (i.e., Th1, Th2, Th17). (Olatunde, Hale, and Lamb 2021) Using recently described scRNA-seq-based cell state signatures (Yasumizu et al. 2024), we examined the Th1, Th2 and Th17-like state of CM CD4 T cells with age. From these analyses, we found that older adults exhibited a significant skewing towards a Th2-like state compared to young adults in CM CD4 T cells that was stably maintained over time. (**Figure 6D, 6E**) Moreover, our follow-up cohort confirmed a continuous increase in the Th2-like state in CM CD4 T cells across age. (**Figure 6F**) Neither Th1- or Th17-like states changed significantly with age or over time in the CM CD4 T cells. To further confirm the association between age and the development of Th2-like state, we utilized our previously published tri-modal TEA-seq data to interrogate TF activity within CM CD4 T cells from children (11-13 yrs) and older adults (55-65 yrs). (**Figure S5H**) (Thomson et al. 2023) Consistent with the elevation in a Th2-like transcriptional state, activity of the Th2-associated TF GATA3 (p=0.00016) was significantly higher in CM CD4 T cells in older adults (**Figure 6G**). The activity of TBX21, the main Th1-associated TF, did not change with age. GATA3 activity also directly correlated with the RNA-based Th2-like cell state metric within this independent dataset. (**Figure 6H**) Consistent with higher GATA3 activity and a Th2-like state, older adults also exhibited increased chromatin openness within the IL4 locus, a GATA3-regulated gene. (**Figure 6I**) Activity of the classical Th2 cytokine-driven TF STAT6 was not elevated in CM CD4 T cell with age (**Figure S5I**), nor was there increased openness at the STAT6-regulated gene IL4R. (**Figure S5J**) The elevation in GATA3 activity in older adult CM CD4 T cells was in the absence of any notable increase in protein levels of circulating Th2-polarizing cytokines IL-4, IL-5 and IL-13 across age. (**Figure S5K, S5L**) Thus, we find that the molecular programming of memory CD4 T cells become gradually skewed towards a GATA3-related transcriptional state with age that is not directly reflected on a systemic level in the circulating proteome.

The age-related transcriptional changes in memory T cells collectively indicate the potential for altered B cell activation (via CD40LG:CD40 interactions) and B cell class switching (via Th2 state) in response to vaccine-induced perturbation in older adults. To further link stable homeostatic alterations in memory T cells with broad vaccine-induced memory B cell dysregulation with age, we interrogated the association of Th2 and T follicular helper cell states in CM CD4 T cells with chronological age, transcriptional age (RNA age metrics), baseline T cell functionality and broad vaccine-induced B cell responses in our longitudinal cohort. (**Figure 7A**) Consistent with the direction of gene expression, Th2 and Tfh states correlated with up- and down-summary metrics respectively (Th2:RNA Age metric (up) pval=0.02, rho=0.28; Tfh:RNA Age metric (down) pval=1.02e-10, rho=0.796). Th2 and Tfh states were commonly associated with many features of T cell functionality and B cell responses (albeit in opposite directions), including CM CD4 T cell CXCR5 expression and core memory B cell activation pathways. Notably, the Th2 state of CM CD4 T cells specifically and inversely correlated with CD40L:CD40 interactions between CM CD4 T cells and core memory B cells (confirmed in our follow-up cohort (pval=0.011, rho=-0.17)), as well as the magnitude of change in HAI inhibition (i.e., functional antibody responses) at day 7 post-vaccination. (**Figure 7B**) Additional, we find IGHG4 expression in Core memory and CD95 memory B cell post-vaccination positively correlated with Th2 state of CM CD4 T cells but not Tfh programming (**Figure 7A, 7B**), which is consistent with cytokines mediating changes in B cell antibody isotype. These data build a model in which memory CD4 T cells progressively lose intrinsic helper potential in tandem with developing a Th2 state across age that leads to reduced memory B cell activation, altered class-switching and less function antibody production in older adults. Taken together, exploration of our new, large-scale longitudinal immune health resource reveals novel cellular and molecular features of the healthy immune system connected to its responsiveness to acute and chronic perturbation with age.

**Figure 7.**
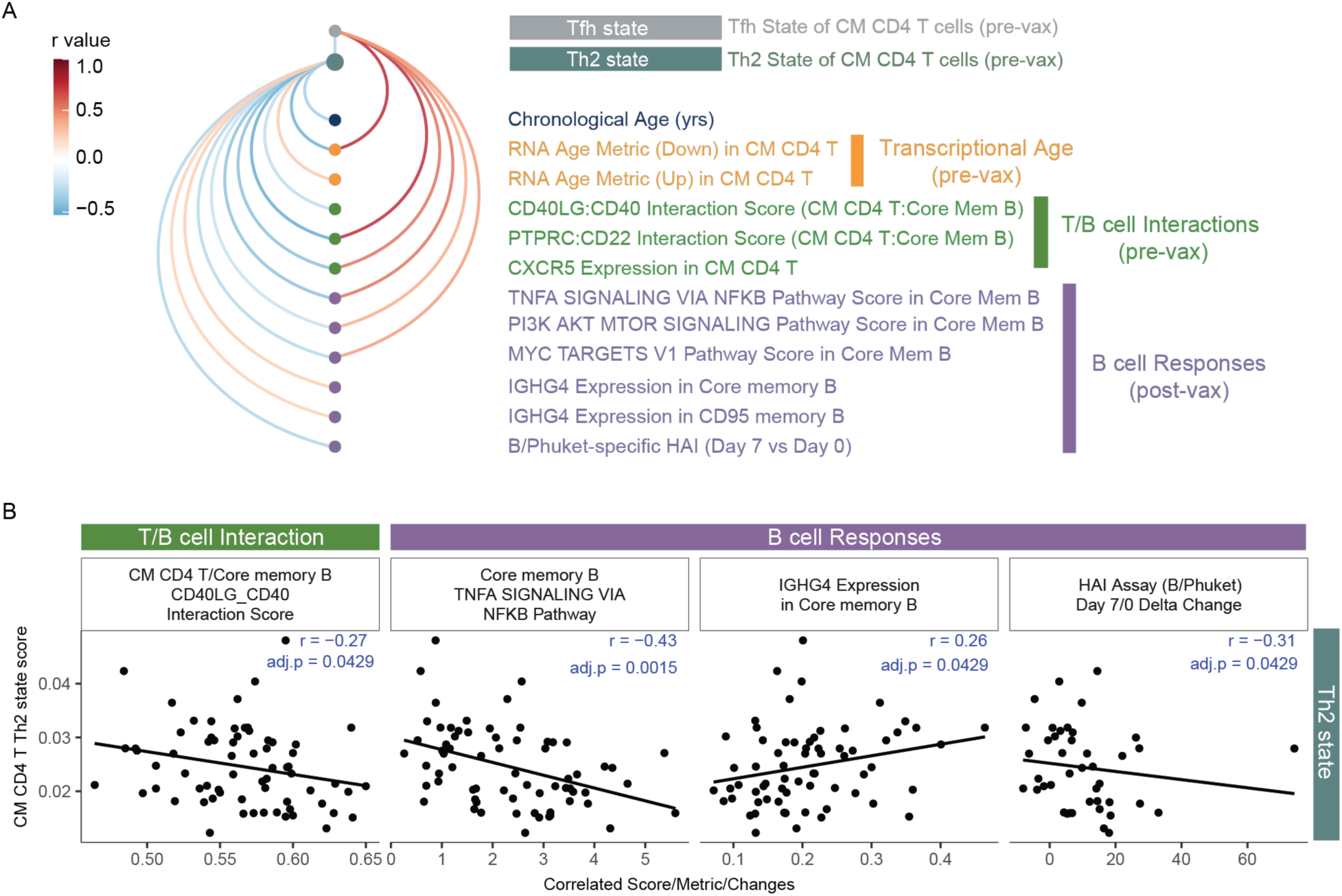
Age-related transcriptional states of central memory CD4 T associated with memory B cell response to influenza vaccination. **A.** Arc plot of Th2 and Tfh cell states in CM CD4 T cells correlations with features of age, T cell - B cell interactions and B cell responses to flu vaccination. Only correlations with pval <0.05 are shown. **B.** Select Spearman correlations of Th2 state with T-B interactions and B cell responses.

## Discussion

In this study, we sought to uncover key features of the healthy peripheral immune system that accompany the aging process. By applying our newly developed Human Immune Health Atlas to two unique healthy adult cohorts (one longitudinal and one cross-sectional), we conducted an in-depth analysis of the molecular and cellular landscape of the immune compartment at homeostasis and in response to acute and chronic perturbation. This effort resulted in a high-resolution scRNA-seq resource comprising over 16 million peripheral immune cells from more than 300 healthy individuals aged 25 to 90 years. Additionally, this resource includes paired plasma proteomics and spectral flow cytometry, offering complex, multi-dimensional insights into human immunity across health and aging. We also provide a suite of interactive data exploration tools at https://apps.allenimmunology.org/aifi/insights/dynamics-imm-health-age/ to enable researchers easier, open access to this extensive human immune health resource and to further facilitate new immunological discoveries and applications.

The longitudinal nature of this study allows us to gain insight into immune cell programming at baseline, its stability over time and its broad alterations with acute and chronic perturbation. Remarkably, we observe long-term stability in homeostatic programming of many immune cell subsets is unaffected by age. While the programming of T cell subsets early in their differentiation was significantly altered by age, there was also little to no age-related effects on the more terminal T cell subsets, contrasting inflammatory, senescent or exhausted T cell features commonly associated in advanced aging studies. (Zhu et al. 2023; Mogilenko et al. 2021; Elyahu et al. 2019) Recent mouse studies have demonstrated a gradual shift towards an exhausted-like transcriptional state within the memory T cell compartment over a decade of antigenic re-challenge. (Soerens et al. 2023) However, our study’s finding on the stability of certain features of transcriptional programming up to 65 years of age suggest this transition towards exhaustion may occur in more advanced age (i.e., >65 yrs of age). It also implies that homeostatic maintenance during early adulthood may be a more nuanced relationship between a cell’s lifespan, its programming, and the microenvironment it inhabits (i.e., lymph nodes) in the absence of overt disease. (Ural et al. 2022; Cakala-Jakimowicz, Kolodziej-Wojnar, and Puzianowska-Kuznicka 2021)

CMV infection is another immunologic challenge intertwined with age - implicated to cause chronic immune activation and to modulate protective immunity in adults. (Furman et al. 2015; Picarda and Benedict 2018) Similar to previous observations, we found that individuals of all ages infected with CMV had a marked shift in their peripheral immune landscape, highlighted by the acquisition of terminally differentiated cytotoxic T cell subsets and adaptive NK cells. (Bayard et al. 2016; Wertheimer et al. 2014; Schlub et al. 2011) Moreover, these changes were stable over the 2-year course of the study, neither displaying transient nor cumulative reprogramming as one may expect from reoccurring or continuous immune perturbation. Adaptive NK cells show an age-associated decrease in frequencies and subtle transcriptional differences suggestive of reduced activation in older adults. The gradual age-related reduction in ZBTB38 and KLRC2 expression, key population-defining markers recently described for ‘NK3’ cells (Rebuffet et al. 2024), also indicates potential plasticity in the epigenetic programming of this immune cell subset across age. (Lau et al. 2018; Rückert et al. 2022; Schlums et al. 2015) Of note, the recruitment of donors for this study occurred during the COVID pandemic, a period when exposure to common viruses was significantly reduced, and thus, the stability of the CMV-induced immune landscape with age in our study may uniquely reflect less immune remodeling by specific and bystander viral exposure during this period. (Nikolich-Žugich et al. 2020) Given that CMV activation and exposure significantly impact the success of solid-organ transplantation (Raglow and Kaul 2023; Kotton 2013; Fishman 2007), further research into the persistent cytotoxic reshaping of the immune cell landscape and its potential role in immune-mediated disease risk is warranted.

In addition to studying the impact of age on the immune landscape during CMV-mediated perturbation, we deeply examined acute, age-specific transcriptional responses of memory B cells to vaccine-induced immune perturbation. A unique feature of our Immune Health Atlas is the detailed annotation of the B cell compartment, which revealed novel age-specific transcriptional states in effector and memory B cell subsets in tandem with changes in vaccine-specific antibody responses during multiple rounds of immune activation. We found altered transcriptional states post-vaccination in plasma cells and, interestingly, effector memory B cells, a unique B cell subset associated with both age and viral infection, from older adults. This CD27^−^CD11c^+^Tbet^+^Zeb2^+^ effector memory B cell subset described had age-associated transcriptional state that corresponded to reduced activation potential via lower ROS pathway activity and lower frequencies of IgG^+^ cells, highlighting a potential mechanism for reduced protective responses with age. (Burton et al. 2022). These effector B cells in older adults also exhibited a reduced gene program affiliated with long-lived humoral responses in the context of influenza vaccination. (Nellore et al. 2023) Thus, our study reveals a new link between effector B cells, protective immunity and aging that may contribute to reduced vaccine efficacy in older adults.

Previously, the connection between reduced B cell responses and age-related transcriptional changes in T cells remained unclear. While evidence suggests CD4 T cells undergo dysregulation with age, findings regarding the directionality and specific features of these age-related changes (e.g., Th1 versus Th2 polarization, effector vs memory function) are conflicting. (Terekhova et al. 2023; Elyahu et al. 2019; Hu et al. 2019) Our findings on the age-related increase in a Th2-like state in central memory CD4 T cells aligns with recent literature finding elevated CCR4+ memory T cells in advanced age (Terekhova et al. 2023) and suggests a mechanism where heightened Th2 programming impedes T-bet activation in older adults. (Dai et al. 2024; Naradikian et al. 2016) Indeed, the only described T-bet deficient patient has been shown to have both high Th2 T cell populations (Yang et al. 2021) and impaired effector B cell development (Yang et al. 2022), directly linking two of our novel observations in this study. Furthermore, efficient generation of effector B cells often relies on contact-dependent help from Tfh cells via CD40 ligand, essential for B cell activation and class switching. (Nonoyama et al. 1993; Song et al. 2022) This further aligns with an elevated Th2 state and reduced CD40L expression in central memory CD4 T cells in older adults. Thus, our study integrates novel mechanisms with established aspects of immune aging, highlighting the dynamic accumulation of features including elevated Th2/Gata3 programming, reduced effector/memory B cell activation, and impaired antibody functionality across age.

In summary, we uncover new insights into the dynamics of the peripheral immune landscape across age, chronic infection and vaccine-induced perturbation, underscoring the importance of longitudinal analyses in facilitating a more comprehensive understanding of age-related immune dysregulation. Our findings demonstrate that the healthy *stable* state of immune cell subsets change with age and impacts immune responsiveness, independent from systemic inflammation and chronic infection. These results have translational implications in the context of designing age-specific vaccines, treating diseases utilizing immune cell-based treatments (e.g., CAR T cells) in older adults, and preventing the onset of age-related immunological diseases such as rheumatoid arthritis.

## Methods

### EXPERIMENTAL MODEL AND SUBJECT DETAILS

#### The Sound Life study cohort

Healthy 25- to 35-year-old and 55- to 65-year-old adult donors were prospectively recruited from the greater Seattle, Washington, USA area as part of the Sound Life Project, a protocol (IRB19-045) approved by the Institutional Review Board (IRB) of the Benaroya Research Institute. All adult participants provided informed consent before participation. Donors were excluded from enrollment if they had a history of chronic disease, autoimmune disease, severe allergy, or chronic infection. All blood samples were collected, processed to PBMCs through a Ficoll-based approach and frozen within 4 hours of blood draw. Plasma samples were processed, aliquoted and frozen within 4 hours of blood draw. Basic demographics are provided in **Supplemental Table 1**.

#### Follow-up cohort

234 paired PBMCs and serum samples were retrospectively selected from a cohort of healthy adults, ages 40 years and older, recruited from the greater Palo Alto, California, USA area, under a protocol approved by Stanford University IRB. (Whiting et al. 2015) All participants provided informed consent before participation and these subsequent studies were approved by the Allen Institute IRB. Basic demographics for the selected follow-up cohort are provided in **Supplemental Table 1**.

### METHOD DETAILS

#### scRNAseq via 10x Genomics v3 chemistry

scRNA-seq was performed on PBMCs from the Sound Life cohort as previously described (Genge et al. 2021), using a modified 10x Genomics Chromium 3′ single-cell gene expression assay with Cell Hashing. In brief, PBMCs from the Sound Life cohort were thawed and stained with oligo-tagged antibodies (HTO) allowing for overloading of Chip G (10x Genomics, PN 20000177) wells at 64,000 cells. At cDNA amplification, HTO additive primer was spiked into the cDNA amplification master mix. Following cDNA amplification per the manufacturer’s instructions, HTO and GEX cDNA products were separated using SPRI-Select (Beckman Coulter, PN B23319) bead-based cleanup before carrying forward into separate library indexing reactions. Libraries were sequenced on a NovaSeq S4 200 cycle flow cell at Northwest Genomic Center at the University of Washington (https://nwgc.gs.washington.edu/). Samples were then computationally resolved and quality-checked using in-house pipelines.

#### scRNAseq via 10x Genomics FLEX

PBMCs from our follow-up cohort were thawed per our standard methodology. (Genge et al. 2021) Samples were run in batches of 48 or 64 samples, with mixed age and sex distributions. PBMC bridging controls were included on each batch to allow for cross batch normalization. Viable samples were processed using Chromium Next GEM Single Cell Fixed RNA Sample Preparation Kit (10x Genomics, PN 1000414) according to the 10x Genomics protocol for “Fixation of Cells and Nuclei”. To facilitate high throughput, volumes were scaled from 1 mL to 250 µL for plate-based sample preparation and handling. Probe hybridization was completed according to the user guide “Chromium Fixed RNA Profiling Reagent Kits for Multiplexed samples” using the Chromium Fixed RNA Kit, Human Transcriptome, 4 rxns x 16 BC (10x Genomics, PN 1000476). Up to 2 million fixed cells were hybridized per sample. After 16-24 hours of probe hybridization incubation at 42°C, samples were pooled at equivalent cell numbers with 15 samples and a bridging control per pool. Each single cell suspension pool was loaded onto two wells of Chip Q (10x Genomics, PN 2000518) for GEM generation at an overloaded concentration of 400,000 cells. Pre-Amplification PCR and library construction were performed per the manufacturer’s instructions. Final scRNAseq libraries were sequenced using a NovaSeq X 25B 300 cycle flow cell or NovaSeq S4 200 cycle flow cell, depending on total read requirements at either Clinical Research Sequencing Platform at the Broad Institute (https://broadclinicallabs.org/) or Northwest Genomic Center at the University of Washington (https://nwgc.gs.washington.edu/). Samples were then computationally resolved and quality-checked using in-house pipelines.

#### Olink Explore 1536

Plasma samples from the Sound Life cohort were run on the Olink Explore 1536 platform, which uses paired antibody proximity extension assays (PEA) and a next generation sequencing (NGS) readout to measure the relative expression of 1472 protein analytes per sample. For plate setup, samples were randomized across plates to achieve a balanced distribution of age and sex. Longitudinal samples from the same participant were run on the same plate. Plasma bridging controls (12-40) were included on each of 6 batches and used for cross-batch normalization.

#### Olink Explore 3072

Serum samples were run on the Olink Explore 3072 platform to measure the relative expression of 2943 proteins. Follow-up cohort samples were split across 3 Olink batches with 41 serum bridging controls included across each of the batches for cross-batch normalization. For plate setup, samples were randomized across plates to achieve a balanced distribution of age and sex.

#### HCMV serology

Viral serology testing for Human Cytomegalovirus (HCMV) was performed at the University of Washington’s Clinical Virology Laboratory in the Department of Laboratory Medicine (https://depts.washington.edu/uwviro/). Plasma or serum samples (200 µL) were run through the FDA-approved LIAISON® CMV IgG Assay to qualitatively detect CMV IgG class antibodies. Results were reported for each sample as ‘Positive’ or ‘Negative’ along with a CMV Ab Screen Index Value ranging from <0.20 to >10.00.

#### Adaptive NK Flow Cytometry

One to two million viable PBMCs were plated into wells of a 96-well U-bottom plate. Cells were stained for viability, FC blocked, and then stained with a surface marker antibody cocktail **(Supplemental Table 3)** using BD Brilliant Staining Buffer (BD Biosciences, PN 563794) for 30 minutes at 4°C. The samples were washed, fixed for 60 minutes at room temperature, FC blocked and permeabilized for 10 minutes at room temperature, then stained with an intranuclear antibody staining cocktail for 60 minutes at 4°C using the eBioscience Foxp3/Transcription Factor Staining Buffer Set (Thermofisher, PN 00-5523-00). After staining, the samples were washed and fixed with Phosphate Buffered Saline solution (PBS) containing 1% paraformaldehyde (PFA) for 15 minutes at 4°C. Finally, the samples were washed and resuspended in PBS before acquisition on a BD Symphony (5L) flow cytometer.

#### B-cell Flow Cytometry

PBMCs were rapidly thawed and diluted into 20% Fetal Bovine Serum (FBS) in ATCC-modified RPMI (ThermoFisher Scientific, PN A1049101) and washed with media. Two million viable PBMCs from each donor were seeded into a 96-well V-bottom plate. PBMCs were then incubated with Human TruStain FcX and Purified mouse IgG (BioRad, PN PMP01X) for 10 minutes at room temperature and washed with Wash Buffer (1% FBS in PBS Solution). Cells were incubated with Fixable Viability Stain 510 (BD Biosciences, PN 564406) for 30 minutes at 4°C and then washed. Cells were stained with a filtered antibody cocktail in Wash Buffer for 30 minutes at room temperature (**Supplemental Table 3**) and finally washed with Wash Buffer. Stained PBMCs were immediately resuspended to 10 x10^6^ cells/mL in Wash Buffer and acquired on an Aurora 5L flow cytometer (Cytek Biosciences).

#### Total Influenza-specific antibody serology

The Meso Scale Discovery (MSD) Prototype Influenza 7-plex Serology Assay protocol measures IgG antibodies in human plasma specific for Influenza vaccine hemagglutinin (HA) antigens: A/Brisbane, A/Hong Kong, A/Michigan, A/Victoria, B/Colorado, B/Phuket, and B/Washington. Briefly, MSD 96-Well 10-Spot multi-array plates coated with seven flu HA antigens were blocked, human plasma samples were diluted 10,000-fold, and added along with HA reference standards and controls to the plate. Plates were shaken for two hours at 15°C to 25°C, washed, then anti-human IgG antibodies labeled with electrochemiluminescent (ECL) SULFO-TAG were added. Plates were shaken for one additional hour at 15°C to 25°C, washed, then MSD GOLD Read Buffer B was added, and the plates were read on an MSD SECTOR S600 ECL plate reader. Test samples were quantified in AU/mL referenced against specific HA reference standards.

#### Hemagglutination inhibition Influenza-specific antibody serology

The MSD 96-well hemagglutination inhibition (HAI) 9-plex Assay measures neutralizing antibodies in human plasma that block the binding of labeled red blood cell vesicles to trimeric Influenza HA antigens, specific for the following lineages: A/Brisbane, A/Cambodia, A/Guangdong, A/HongKong, A/Kansas, A/Shanghai, A/Wisconsin, B/Phuket, and B/Washington. Briefly, plasma samples were first treated with enzymes to remove interfering sialic acid residues. MSD 96-Well 10-Spot multi-array plates coated with nine trimeric flu HA antigens were blocked and then pretreated human plasma samples diluted 5,000-fold along with HA reference standards were added to the plate. Plates were shaken for two hours at 15°C to 25°C, then red blood cell vesicles labeled with ECL SULFO-TAG were added. Plates were shaken for another two hours at 15°C to 25°C, washed, then MSD GOLD Read Buffer B was added, and the plates were read on an MSD SECTOR S600 ECL plate reader. Test samples were reported as Percent Inhibition, relative to a no-plasma diluent only control. Positive samples show high percent inhibition whereas negative or low samples show low percent inhibition.

### QUANTIFICATION AND STATISTICAL ANALYSIS

#### Flow Cytometry data analysis and visualization

Adaptive NK and B cell flow cytometry data analysis consisted of traditional hierarchical gating in FlowJo v10.10 Software. For B cell analysis, total live cell gated data, consisting of all live singlet PBMCs within the experiment, was then downloaded and further processed with the R programming language (http://www.r-project.org) and Bioconductor (http://www.bioconductor.org) software. Data was transformed with an inverse hyperbolic sine (asinh) transformation with a cofactor of 220. Each marker was scaled to the 99.9th percentile of expression of all cells in the experiment. Total live B cells were over-clustered into 100 clusters using FlowSOM with all informative surface molecules as input. Clusters were then hierarchically clustered based on expression of B cell surface molecules and isotype and finally manually assigned to cell subsets, as described previously. (Glass et al. 2020) B cell subsets were plotted on a UMAP plot using the umap package in R, based on expression of all informative markers and excluding isotype.

#### Olink data processing

Olink’s standard data normalization was performed on these datasets. Protein expression values were first normalized across wells using an internal extension control (IgG antibodies conjugated with a matching oligo pair). Plates were then standardized by normalizing to the inter-plate pooled serum or plasma controls run in triplicate on each plate. Data were then intensity normalized across all cohort samples. Final normalized relative protein quantities were reported as log2 normalized protein expression (NPX) values.

#### AIFI Immune Health Atlas (V1) building

To build our scRNA-seq PBMCs dataset for the AIFI Immune Health Atlas, we utilized data and analysis environments within the Human Immune System Explorer (HISE) system (https://allenimmunology.org/) to trace data processing, analytical code, and analysis environments from the original, raw FASTQ data to our final, assembled reference atlas. (Meijer et al. 2024) A graph representation of the analysis trace for this project is available at https://apps.allenimmunology.org/aifi/resources/imm-health-atlas/reproducibility/. Additional details are available in our analysis notebooks on Github at https://github.com/aifimmunology/aifi-healthy-pbmc-reference and at our website at https://apps.allenimmunology.org/aifi/resources/imm-health-atlas/.

#### Pipeline processing

After sequencing, Gene Expression and Hash Tag Oligonucleotide libraries from pooled samples in our pipeline batches were demultiplexed and assembled as individual files for each biological sample as described in (Swanson et al. 2022).

#### Data selection from HISE storage

To assemble the input data for our reference dataset, we selected samples from all subjects in our longitudinal cohort (Sound Life cohort; Young Adult, age 25-35 yrs and Older Adult, age 55-65 yrs) and 16 samples from a previously described healthy pediatric cohort collected at the University of Pennsylvania (age 11-13 yrs (Thomson et al. 2023)). In total, 108 samples were selected for use in our reference (Pediatric, n = 16; Young Adult, n = 47; Older Adult, n = 45), consisting of 2,093,078 cells before additional QC filtering.

#### Labeling and doublet detection

To guide cell type identification, we labeled cells from each sample using CellTypist (v1.6.1)(Domínguez Conde et al. 2022), using the following reference models: Immune_All_High (32 cell types), Immune_All_Low (98 cell types), and Healthy_COVID19_PBMC (51 cell types) by following the approach described in the CellTypist reference documentation at https://celltypist.org. We also labeled our cells using Seurat (v5.0.1) (Hao et al. 2021), which was downloaded from the Zenodo repository (DOI: https://zenodo.org/doi/10.5281/zenodo.7779016). For Seurat labeling, we utilized the SCTransform function to transform data to match the reference dataset, then used FindTransferAnchors and MapQuery to assign cell types based on the 3-level cell type labels provided in the reference dataset. We performed doublet detection using the scrublet package (v0.2.3) (Wolock, Lopez, and Klein 2019) implemented in the scanpy.external module of scanpy (v1.9.6).

#### QC filtering

After assembly of all cells across our 108 samples, we filtered our data against a set of QC criteria designed to remove possible doublets (based on scrublet and high gene detection, > 5,000 genes), low-quality cells (based on low gene detection, < 200 genes), and dying or dead cells (based on mitochondrial UMI content, > 10%). Mitochondrial content was assessed using scanpy by identifying mitochondrial genes (prefixed with “MT-”), and using the scanpy function calculate_qc_metrics, as demonstrated in the scanpy tutorials (https://scanpy-tutorials.readthedocs.io/en/latest/pbmc3k.html). In total, 140,950 cells were removed (6.73%), and 1,952,128 cells passed QC filtering (93.27%).

#### Clustering and cell subsetting

After QC filtering, all remaining cells were clustered using a Scanpy workflow (Wolf, Angerer, and Theis 2018) to normalize, log transform, perform PCA, integrate age groups with Harmony (Korsunsky et al. 2019), perform Leiden clustering (Traag, Waltman, and van Eck 2019), and generate 2-dimensional UMAP projections. Clusters were assigned to major cell classes based on marker gene detection. For each cluster, the fraction of cells with detected gene expression for each marker was computed, and clusters meeting a set of cell class-specific criteria were selected for downstream annotation by our domain experts. Clusters from specific cell classes were selected from the full dataset for iterative clustering analysis where necessary to identify cell subpopulations with additional resolution. To assist with cell labeling, marker genes were identified for each cluster at each level of iteration and at each clustering resolution using the scanpy function scanpy.tl.rank_genes_groups with the parameter method = ‘wilcoxon’.

#### Expert annotation of cell types

Following high-level and iterative clustering, teams of domain experts within the Allen Institute for Immunology examined marker gene expression for clustered datasets and assigned cell type identities to each cluster. As part of this cell type identification exercise, any remaining doublets or low-quality clusters were also identified for later removal during dataset assembly. Cell type nomenclature and multiple levels of cell type resolution were harmonized across our domain experts. We identified 9 low-resolution cell classes (AIFI_L1), 29 mid-level cell classes (AIFI_L2), and 71 high-resolution cell classes (AIFI_L3). After annotation, cluster labels were transferred to individual cell barcodes to assemble a final set of labels across all cells in our dataset.

#### Training cell type labeling models

In order to utilize our cell type atlas to label other PBMCs datasets, we used CellTypist (v1.6.2 (Domínguez Conde et al. 2022)) to generate cell type labeling models. CellTypist utilizes logistic regression as part of its model generation process with a One-vs-Rest (OvR) approach to multiple classes. We slightly modified CellTypist v1.6.2 to also allow the multinomial method provided by the LogisticRegression() function in the scikit.learn Python package, available at https://github.com/aifimmunology/multicelltypist. We used this modified version of CellTypist to train models for each level of our cell type labels using these two approaches. We utilized OvR regression for AIFI_L1 and AIFI_L2 labels, and Multinomial regression for AIFI_L3.

#### Sound Life scRNA-seq dataset assembly

To build our longitudinally sampled PBMCs scRNA-seq dataset, we utilized data and analysis environments within the HISE system to trace data processing, analytical code, and analysis environments from raw FASTQ data to a labeled, high-quality final dataset. (Meijer et al. 2024) A graph representation of the analysis trace for this project is available at https://apps.allenimmunology.org/aifi/insights/dynamics-imm-health-age/reproducibility/. Additional details are available in our analysis notebooks on Github https://github.com/aifimmunology/sound-life-scrna-analysis/ and at our website https://apps.allenimmunology.org/aifi/insights/dynamics-imm-health-age/.

#### Pipeline processing

After sequencing, Gene Expression and Hash Tag Oligonucleotide libraries from pooled samples in our pipeline batches were demultiplexed and assembled as individual files for each biological sample, as described above for the PBMCs Atlas dataset.

#### Data Selection

Input data for our dataset was assembled as described above for the PBMCs Atlas dataset. In total, 868 samples were selected for use in our dataset (Young Adult, n = 49 subjects; n = 418 samples; Older Adult, n = 47 subjects, n = 450 samples), consisting of 15,781,886 cells before additional QC filtering.

#### Labeling and doublet detection

To add cell type labels, we utilized CellTypist (v1.6.1) (Domínguez Conde et al. 2022) and the CellTypist models we generated using our Immune Health Atlas dataset at three levels of cell type resolution (AIFI_L1, 9 broad cell classes; AIFI_L2, 29 intermediate cell classes, and AIFI_L3, 71 high-resolution cell classes) by following the approach described in the CellTypist reference documentation at https://celltypist.org. We performed doublet detection the same way as described above for the Atlas. After cell type labeling and doublet detection were performed on each sample, we assembled data, labels, scrublet calls, sample metadata, CMV status, and BMI data across eight sets of samples defined by the cohort, biological sex, and CMV status of subjects to facilitate downstream analyses.

#### QC filtering

After assembly of all cells across our samples, we filtered our data against the same set of QC criteria as described above for the PBMC Atlas. QC criteria were applied to the dataset in series: cells retained were not identified as a doublet by Scrublet; the percent of total UMIs assigned to mitochondrial regions was < 10%; N genes detected per cell was > 200 and < 2,500. In total, 951,864 cells were removed (6.03%), and 14,830,022 cells passed QC filtering (93.97%).

#### Clustering, subsetting, and doublet removal

After QC filtering, all remaining cells were clustered within AIFI_L2 labels using the scanpy workflow described above for the PBMC Atlas. lusters were filtered to remove doublets based on marker gene expression. For example, CD4 naive T cell clusters with a high fraction of MS4A1 expression were removed as B cell doublets. We also removed clusters with low gene detection as low quality. Due to their generally lower gene detection, this threshold was not applied to Erythrocyte or Platelet populations. In total, 800,059 cells (5.39%) were removed. 14,029,963 cells (94.61%) were retained for final clean up based on high-resolution AIFI_L3 labels.

#### Refinement of cell type labels

After removal of doublets at the AIFI_L2 cell type resolution, we assembled cells from all samples for each cell type in the high-resolution AIFI_L3 labeling results. Separately for each cell type, we performed the scanpy data processing procedure described above to enable a final review of cell type labels. Each cell type was examined to identify remaining doublet clusters or clusters of cells that appeared to be mislabeled based on marker gene expression in collaboration with immunological domain experts. During this process, 234,117 cells were removed. 13,795,846 cells were retained for the final reference dataset and downstream analyses.

#### Follow-up Cohort scRNA-seq Reference

Using our Atlas dataset, all cells in the follow-up cohort were automatically annotated with custom label transfer models built using the CellTypist framework. First, the genes in the reference dataset were subsampled to match the 18,000 genes in the 10x FLEX scRNA-seq assay. This improved model performance by enhancing recognition of more rare cell populations. Then, three models were built for label transferring – one for each cell labeling level. Finally, the follow-up cohort data was labeled. Dataset cleanup, quality checking, clustering, and visualization were carried out using Scanpy. (Wolf, Angerer, and Theis 2018) Doublets were detected and removed using Scrublet, and minimal inter-batch effects were adjusted in the PCA space for visualization purposes using Harmony. (Korsunsky et al. 2019) After QC filtering, labeling and data clean-up, 3,627,005 cells were retained in the final follow-up reference dataset and used for downstream analyses.

#### Olink Analysis

The Olink analysis was performed in R using the lme4 package (Bates et al. 2014), applying the design formula: NPX(Bridged) ~ Age Group + CMV + Sex. The comparisons were based on the age group factor. Proteins with an adjusted p-value below 0.05 were considered differentially expressed.

#### Cell frequency comparison from scRNA-seq

To deal with the constant sum issue on compositional data, we applied the centered log ratio (CLR) transformation on the frequency data for each sample. This transformation was applied to 71 cell types at level 3 labels for each scRNA sample in most frequency comparisons. For comparing switched and non-switched CD27-effector B cell in scRNA data, the CLR transformation was based on total memory B cells at level 3 labels. For the Tfh flow data comparison, the transformation was based on total T cells. In the B cell flow data comparison, the CLR transformation included 9 B cell types and 1 non-B cell type. For the isotype in the B cell flow data experiment, CLR transformation was performed on individual cell types for different isotype. For paired data, we applied the signed-rank Wilcoxon test to the CLR-transformed data. For unpaired data, we used the rank-sum Wilcoxon test on the CLR-transformed data.

#### DEG analysis

Differential gene analysis was performed using the DESeq2 (version 1.42.0 (Love, Huber, and Anders 2014)) package in R (version 4.3.2). For all comparisons, genes were filtered based on a minimum of 10% non-zero counts in each comparison.

Aggregated counts from single cells were used for the comparisons. Several DESeq2 analyses were conducted with default setting:

- For the global baseline comparison of flu year 1 day 0 samples, the design formula used was: Aggregated Counts ~ Age Group + CMV + Sex. The contrast was made for each factor.
- For the global flu vaccine and no-vaccine comparisons of year 1 and year 2 samples, the design formula used was: Aggregated Counts ~ Visit + Age Group+ CMV + Sex + Subject. The contrast was made for the visit factor (day 0 vs day 7).
- For the flu response comparison between two age groups for aligned flu years, we included only donors with samples from both the 2020-2021 and 2021-2022 flu seasons. Day 7 samples from both years were used in the analysis. The design formula applied was: Aggregated Counts ~ Flu Year+ Age Group+ CMV + Sex. The contrast was made on the age group factor (young vs older).
- For the CMV comparison, we performed DESeq2 analysis using the design formula: Aggregated Counts ~ Age Group+CMV + Sex. The contrast was made for the CMV factor (positive vs negative).

Genes were considered differentially expressed if their adjusted p-value was below 0.05 and their absolute fold change was greater than 0.1.

#### NMF projection

We used the NMF projection tool (https://github.com/yyoshiaki/NMFprojection/tree/main) with its precomputed NMF matrix (https://github.com/yyoshiaki/NMFprojection/blob/main/data/NMF.W.CD4T.csv.gz) to compute the NMF scores on Python (version 3.10.13). For each sample, we first subsetted the CD4 T cells and then performed NMF projection according to the recommended workflow. We normalized the different NMF scores based on the mean scores across cells in each cell type at the sample level. For the top genes defining each NMF factor, we extracted the weight of each gene from the precomputed NMF matrix. Genes with higher weights were considered to be more contributive to each factor. We choose the top 20 genes for Tfh NMF factor with highest weight.

#### Cell to cell interaction analysis

We conducted cell to cell interaction analysis by using CellphoneDB (v5 https://github.com/ventolab/CellphoneDB) with method: statistical inference of interaction specificity on default setting. For each sample, we calculated interactions for CM CD4 T cells and core memory B cells. We retained the interactions with an adjusted p-value below 0.05. For group comparisons, we applied a threshold of n>15 interaction numbers in two groups (young or older) to filter out the interaction with small numbers of significance.

#### Enrichment analysis

Two types of enrichment analysis were conducted. For Gene Set Enrichment Analysis (GSEA), using the results from DESeq2, we ranked the genes based on −log10(p-value) * sign(log2FoldChange) and performed enrichment analysis using the fgsea R package (version 1.28.0). Pathways with an adjusted p-value lower than 0.05 were kept. For Sample Level Enrichment Analysis (SLEA), we used genes that passed the 10% minimal expression threshold as the background gene set and calculated pathway scores for each sample using the leading-edge genes from the fgsea results. At the pseudobulk level, we calculated the mean gene expression with random genes from the background set for each sample, performing 1000 iterations. The z-score was computed as the deviation between the observed mean of leading-edge genes and the mean of the permuted means of random genes, divided by the standard deviation of the permuted means. This deviation was used as the pathway score for each sample.

#### TEA-seq analysis

TEA-seq datasets were directly downloaded from GEO (https://www.ncbi.nlm.nih.gov/geo/query/acc.cgi?acc=GSE214546). We performed label transfer, doublet detection, and quality control (QC) based on the RNA module. Label transfer was conducted using Celltypist, based on the models generated from our atlas reference at three levels. We filtered the cells if predicted labels are not T cells. Doublet detection was done using the scanpy.external.pp.scrublet function on the RNA module only. Predicted doublets were filtered out before any downstream processes. QC was performed by ensuring that pct_counts_mito was below 15%, and n_genes_by_counts was either below 2500 or above 200. Cells from the RNA module were matched to those from the ATAC module using original barcodes and well IDs. Any mismatched cells containing only RNA or only ATAC modules were excluded. We performed scATAC analysis on ArchR (version 1.0.2). (Granja et al. 2021) The following steps were executed using the default ArchR workflow: LSI dimensionality reduction, group coverage, identification of reproducible peaks (MACS3) (Y. Zhang et al. 2008), peak matrix construction, motif annotation (cisbp database), background peak construction, deviation matrix, and weight imputation (MAGIC). (van Dijk et al. 2018) The z-scored ChromVar motif matrix was extracted with imputed weights. The motif scores at the sample level were calculated based on the mean score for each cell type within each sample.FOli

### RESOURCE AVAILABILITY

#### Lead contact

Further information and requests for resources and reagents should be directed to and will be fulfilled by lead contacts, Claire E. Gustafson and Peter J. Skene.

#### Materials availability

This study did not generate new unique reagents.

#### Data and code availability

Single-cell RNA-seq data will be deposited at GEO (*to be released upon publication*). Raw single-cell RNA-seq fastq files will be deposited at dbGap and be publicly available as of the date of publication. All original code has been deposited at GitHub and Zenodo: https://github.com/aifimmunology/aifi-healthy-pbmc-reference/ (https://zenodo.org/records/13352146) for generation of the Immune Health Atlas, https://github.com/aifimmunology/sound-life-scrna-analysis/ (https://zenodo.org/records/13352142) for labeling and assembly of the Sound Life Cohort data, and https://github.com/aifimmunology/IHA-Figure (*Zenodo link will be publicly available as of the date of publication*) for downstream analysis and figure generation. All processed data, data objects, and clinical metadata derived from de-identified human subjects is available at the Human Immune System Explorer (HISE) website. This paper also analyzes existing, publicly available data (GEO accession: GSE214546). Any additional information required to reanalyze the data reported in this paper is available from the lead contact upon request.

#### Data Visualizations

The following tools are provided to facilitate data exploration and discovery:

**Human Immune Health Atlas**: https://apps.allenimmunology.org/aifi/resources/imm-health-atlas/

1. ***AIFI Immune Health Atlas UMAP Explorer***. Explore cell subsets and gene expression in a UMAP viewer to display an overview of the entire dataset. https://apps.allenimmunology.org/aifi/resources/imm-health-atlas/vis/umap/
2. ***AIFI Immune Health Atlas Gene Expression Reference***. A quick reference for gene expression across cell types and age groups. https://apps.allenimmunology.org/aifi/resources/imm-health-atlas/vis/reference/
3. ***AIFI Immune Health Atlas Clinical Data Explorer***. Flexible visualization of clinical metadata and lab results collected from our Atlas subjects. https://apps.allenimmunology.org/aifi/resources/imm-health-atlas/vis/clinical/

**Dynamics of Human Immune Health and Age Resource:** https://apps.allenimmunology.org/aifi/insights/dynamics-imm-health-age/

4) ***Sound Life Longitudinal Dynamics Explorer***. Explore gene expression and cell type frequency across the longitudinally sampled blood draws within each healthy subject of our longitudinal cohort. https://apps.allenimmunology.org/aifi/insights/dynamics-imm-health-age/vis/dynamics/
5) ***Sound Life Differential Gene Explorer***. Browse results of pairwise differentially expressed gene (DEG) tests for age, sex, CMV and influenza vaccination between our cohort samples. https://apps.allenimmunology.org/aifi/insights/dynamics-imm-health-age/vis/deg/
6) ***Sound Life Clinical Data Explorer***. Flexible visualization of clinical metadata and lab results collected from our cohort’s subjects. https://apps.allenimmunology.org/aifi/insights/dynamics-imm-health-age/vis/clinical/

## Acknowledgments

We thank the study participants and the clinical research team at Benaroya Research Institute and Stanford University, including H. Maecker and E.Krishnan, for their effort and dedication to this project research. We thank the Allen Institute founder, P.G. Allen, for his vision, encouragement and support. We also thank H.Gustafson (https://earlyfutures.com/) for her collaboration on the Immune Health Atlas color palette and all members of the Allen Institute for Immunology, in particular the facilities and operations teams who helped maintain the productive research environment and the Human Immune System Explorer (HISE) software development team for their constant support. Research reported in this publication was supported by the Allen Institute, Benaroya Research Institute and by National Institute on Aging award K01AG068373 (C.E.G.). The content is solely the responsibility of the authors and does not necessarily represent the official views of the National Institutes of Health. Overview figures were created with BioRender.com.

---

**Supplemental Figure 1, in regard to Main Figure 2.**
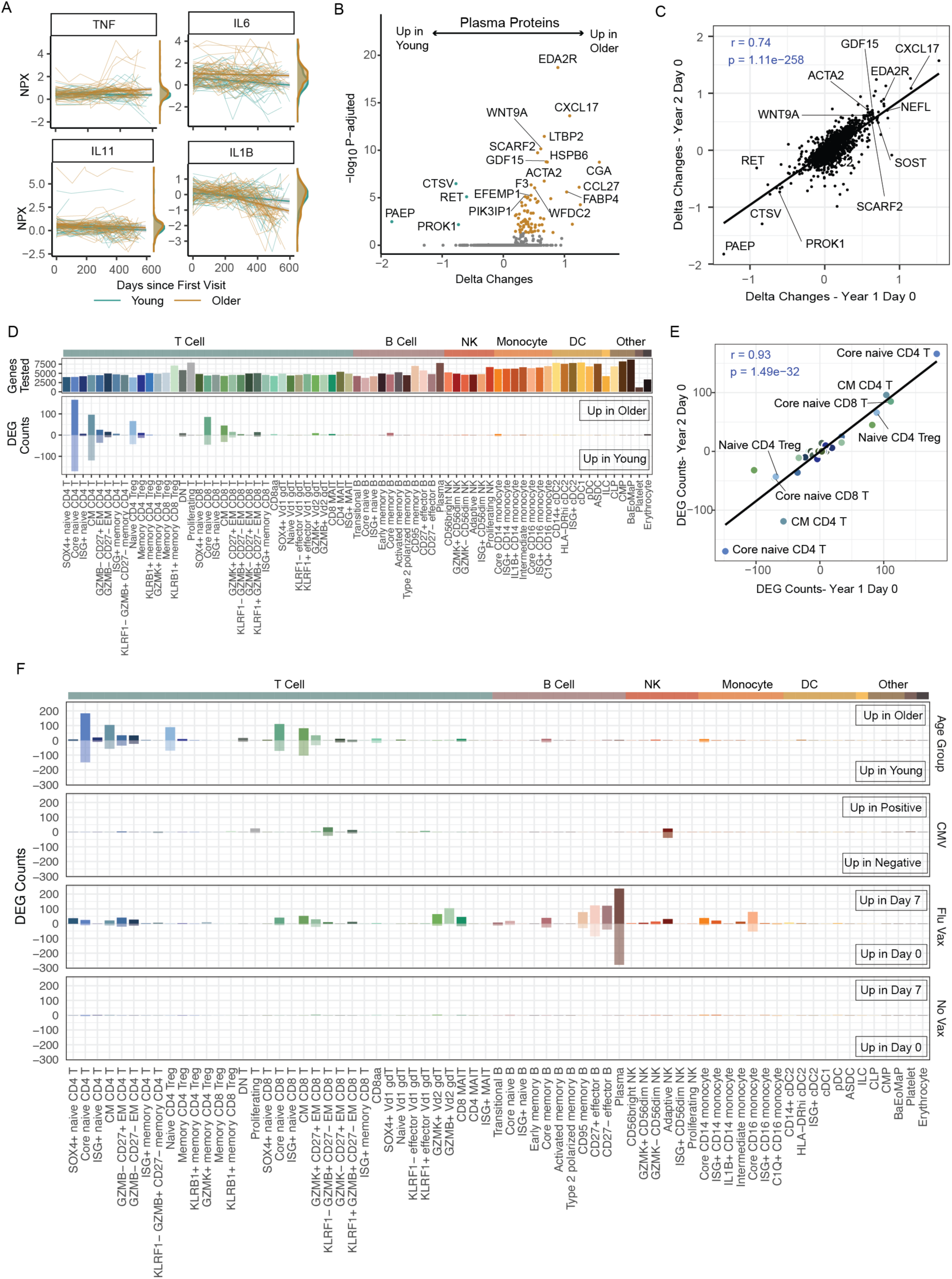
Impact of age, CMV infection and influenza vaccination on the healthy immune landscape. **A.** Normalized protein expression (NPX) of inflammatory markers TNF, IL-6, IL-1b and IL-11 over time in young (teal) and older (bronze) adult plasma. **B**. Volcano plot of the age-related expression differences in circulating plasma proteome at baseline (Flu Vax Year 2 Day 0). **C**. Spearman correlation between age-related protein expression difference at Year 1 and Year 2. **D.** The number of down-regulated or up-regulated differential expressed genes (DEGs) from DEseq2 analysis of immune cell subsets from young (n=40) and older (n=44) adults at ‘Flu Vax year 2 day 0’ using DEseq2 analysis. **E**. Spearman correlation between the number of age-related DEGs in each immune cell subsets in ‘Flu Vax year 1 day 0 and ‘Flu Vax year 2 day 0’. **F.** The number of DEGs in immune cell subsets, comparing CMV (CMV-positive versus CMV-negative), Flu vaccination time series (“Flu Vax”, Day 0 and Day 7 post-flu vaccination) and No Vax time series (“No Vax”, Day 0 vs Day 7 after no vaccination) to that of age. DEGs were defined as log2fc >0.1 and padj<0.05 for all comparisons.

**Supplemental Figure 2, in regard to Main Figure 2.**
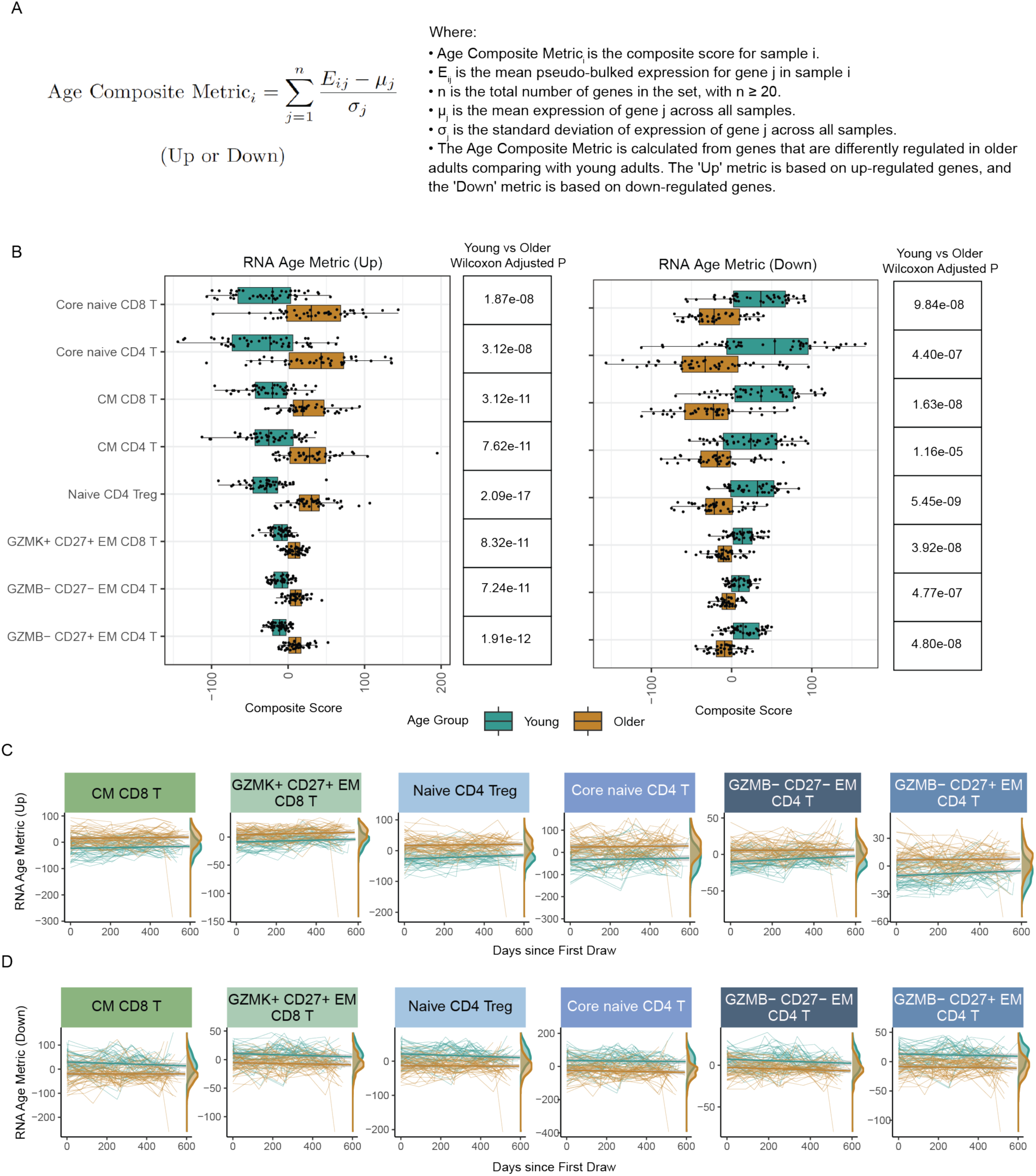
Maintained, age-related transcriptional signatures in healthy immune cell subsets. **A.** Equation for calculating the composite age score in each immune cell subset with more than 20 DEGs between young and older adults. **B.** RNA age metric (upregulated and downregulated genes) in 8 subsets (all subsets with >20 DEGs) comparing young and older adults on year 1 day 0 samples. Each dot is from a single donor. P-value was determined using the Wilcoxon rank-sum test. **C**. RNA age metric (upregulated genes) in select subsets over time in young and older adults. Each donors’ samples are connected with a thin line. **D**. RNA age metric (downregulated genes) in select subsets over time in young and older adults. Each donors’ samples are connected with a thin line.

**Supplemental Figure 3, in regard to Main Figure 4.**
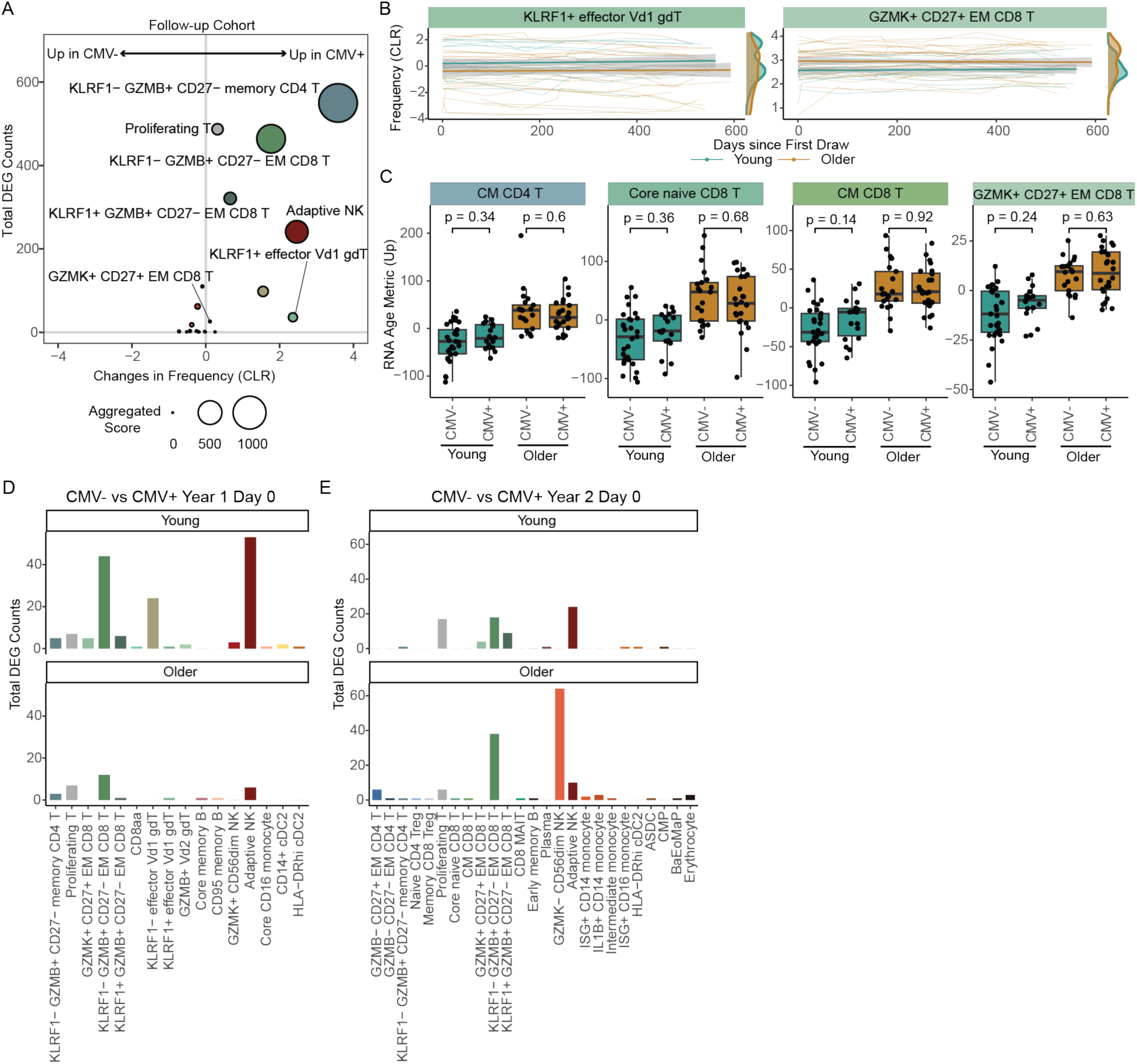
Transcriptional landscape of healthy immune cell subsets altered by CMV and age. **A.** Bubble plot comparison of the change in frequency (using centered log-ratio (CLR) transformation) and number of DEGs between CMV+ (n=136) and CMV- (n=97) adults in our follow-up cohort. Bubble size shows a combined metric of change defined as −log10(p.adj from CLR freq comparison) x DEG_Counts**. B.** Select subset frequencies in PBMCs shown over time. Teal dots are young adults. Bronze dots are older adults. Regression line shown. **C.** RNA age metric (upregulated genes) in select T cell subsets split by age and CMV infection status. P-values were calculated using Wilcoxon rank sum test. **D-E.** The number of DEGs (log2fc >0.1 and p.adj<0.05) in immune cell subsets comparing CMV+ and CMV-individuals, in young and older adults separately from our longitudinal cohort at **D.** ‘Flu Vax year 1 day 0’ and **E.** ‘Flu Vax year 2 day 0’.

**Supplemental Figure 4, in regard to Main Figure 5.**
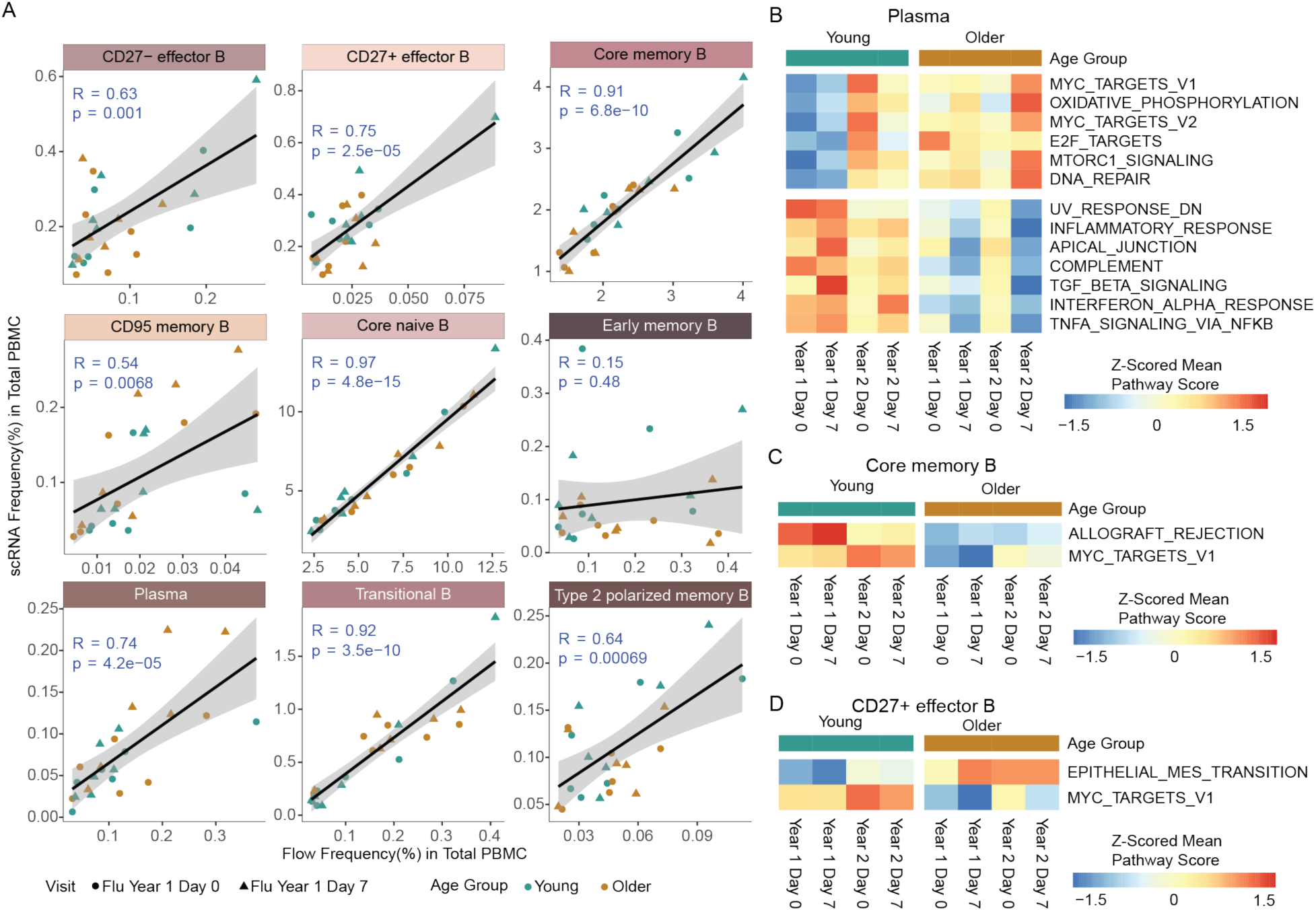
Age-associated responses of non-naive B cell subsets to flu vaccination. **A**. Correlation plots of RNA-quantified (y-axis) and flow cytometry-quantified (x-axis) level 3 B cell population (individual plots) frequencies of total live PBMC. Data shown for 6 young and 6 older adult subjects in flu year 1 that were represented in both scRNA-seq and cytometry analyses. P-value and r values of the Pearson correlation are displayed. **B-D**. Sample level enrichment analysis (SLEA) scores for the top pathways in the Hallmark database at each timepoint for **B.** plasma cells, **C**. core memory B cells and **D**. CD27+ effector B cells in young and older adults.

**Supplemental Figure 5, in regard to Main Figure 6.**
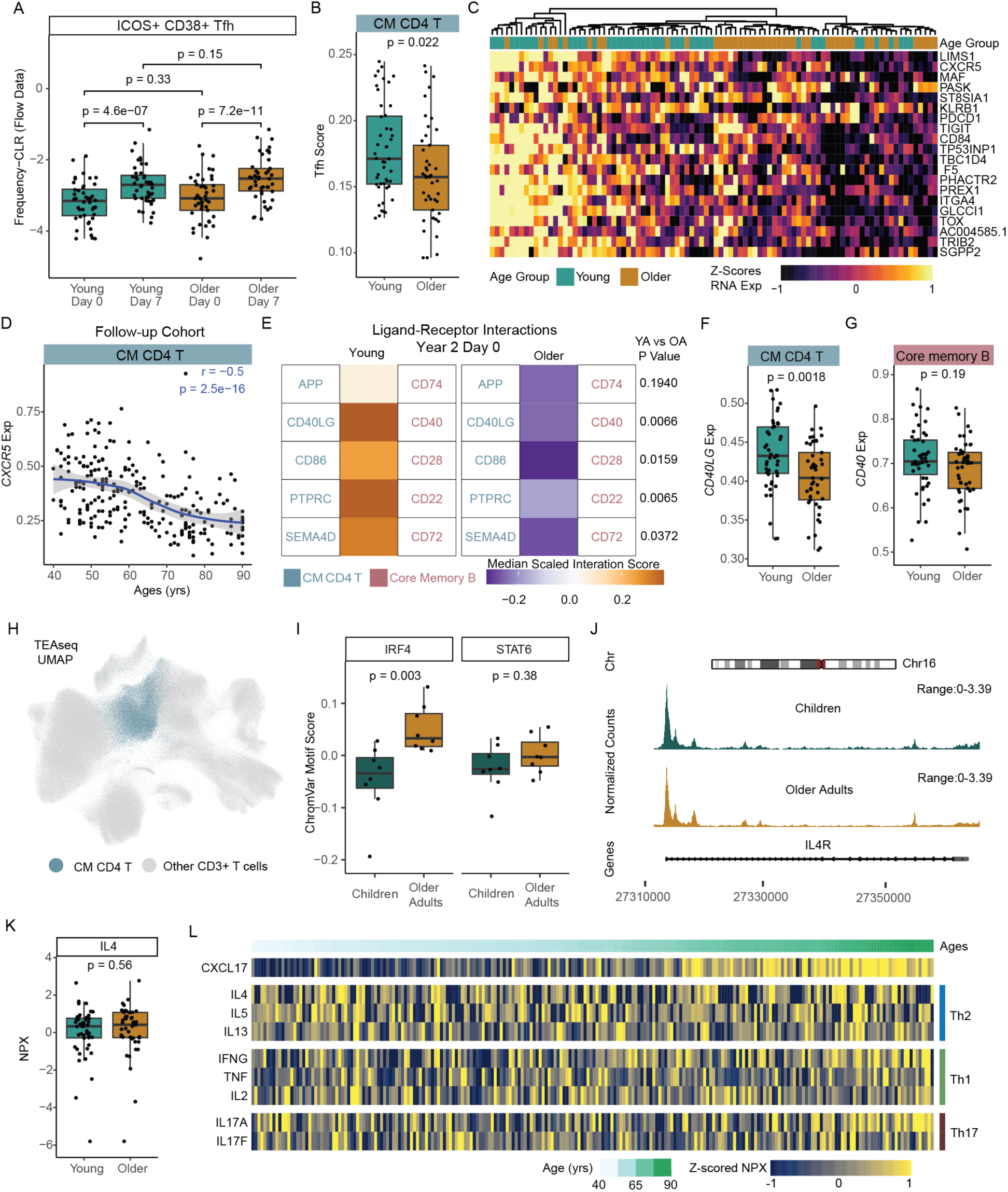
Age-related transcriptional alterations in CM CD4 T cells independent from circulating cytokine signatures. **A.** The CLR transformed frequency of ICOS+ CD38+ Tfh cells determined by spectral flow cytometry, at day 0 and day 7 post-influenza vaccination, comparing responses in young and older adults. P-values were calculated using Wilcoxon’s signed-rank test (paired) for the comparison between Day 0 and Day 7, and using the Wilcoxon rank-sum test for all other comparisons. **B**. Tfh activity score was determined by NMF projection in CM CD4 T cells in young and older adults. **C.** Expression of leading-edge genes in Tfh activity score in young and older adults. **D.** CXCR5 expression in CM CD4 T cells across age in our follow-up cohort. **E.** Receptor-ligand interaction prediction between CM CD4 T cells and core memory B cells in young (n=40) and older (n=44) adults from a single time point (Flu Year 2 Day 0). **F.** CD40LG expression in CM CD4 T cells in young and older adults. **G.** CD40 expression in core memory B cells in young and older adults. **H.** TEAseq UMAP of T cells based on RNA module, with CM CD4 T cells highlighted. **I**. IRF4 and STAT6 transcription factor (TF) activity based on Chromvar analysis of scATACseq data in CM CD4 T cells from children (n=8) and older adults (n=8). P-value was determined by Wilcoxon rank-sum or Wilcoxon signed-rank test, as appropriate, unless otherwise indicated in legend. **J.** Chromatin accessibility tracks of the *IL4R* gene region in CM CD4 T cell subsets from TEA-seq data, showing normalized read coverage. **K**. IL-4 normalized protein expression (NPX) in young (teal) and older (bronze) adult plasma. **L**. Normalized protein expression (NPX) of select Th1-, Th2- and Th17-related serum proteins in our follow-up cohort, with donors ordered by age.

