## Supplemental Methods for "Longitudinal Multi-omic Immune Profiling Reveals Age-Related Immune Cell Dynamics in Healthy Adults"

#### T-cell flow cytometry

2 million PBMCs were added to wells of a 96-well plate and centrifuged at 750g for 5 minutes at 4◦C. Cell pellets were resuspended to a final volume of 100 µL with a viability master mix containing 1:400 FVS510 (BD, 1 μg/mL) and 1:50 Human TruStain FcX Fc Receptor Blocking Solution (BioLegend, PN 422302) in PBS, and incubated for 30 minutes at 4°C. After one wash with Cell Staining Buffer (BioLegend, PN 420201), the samples were stained with a surface marker antibody cocktail (**Supplemental Table 3**) and 1:10 Brilliant Buffer Plus (BD, PN 566385) in Cell Staining Buffer for 30 minutes at 4°C. After two washes with Cell Staining Buffer, the samples were resuspended in FluoroFix fixation buffer (BioLegend, PN 422101) and incubated for 30 minutes at room temperature. After an additional wash, the samples were resuspended in Cell Staining buffer and stored at 4°C until acquisition. Samples were acquired within 1 day of storage using an Aurora 5L flow cytometer (Cytek Biosciences).

#### T cell FlowCyto analysis.

For automated cell type identification, following unmixing and compensation, FCS files were first pre-processed with Flowcut package (Meskas, Wang, and Brinkman 2020) to remove technical abnormal signals. Logicle transformation was applied to all fluorescent channels. To remove doublets, a preset adaptive gating template was applied to the data using the OpenCyto package (Finak et al. 2014). Then, an LDA model was trained and applied to remove debris. To facilitate cell type annotation, a machine learning model were developed for based on Cyanno framework (Kaushik et al. 2021). In brief, a subset of flow cytometry data was selected and unsupervised clustered using Flowsom (Van Gassen et al. 2015). The clusters were manually annotated based on phenotype to create a labeled training dataset. Then the model was trained and applied to all samples. Cell frequencies out of live cells and MFI for all markers were extracted for each gated population and used for downstream analysis. CM CD4 T cells were identified in all flu vaccine day 0 and day 7 samples via our Cyanno pipeline and all cells were combined together for downstream processes. We selected the top 15 variable genes out of 25 as input for PCA dimensionality reduction. The data was normalized using scyan.preprocess.scale before PCA. Batch correction was performed using scapy.external.harmony on the batch factor. Leiden clustering was performed on the first 10 harmonized principal components (PCs) with resolutions of 0.25, 0.5, 0.75, 1.5, and 2. The result from the 1.5 resolution was selected. Based on the clustering results, two populations of T follicular helper (Tfh) cells (CXCR5+ PD1- and CXCR5+ PD1+) were identified within central memory (CM) CD4 T cells. We further subset the CXCR5+ PD1+ Tfh cells for additional leiden clustering and identified ICOS+ CD38+ Tfh cells among them at Leiden resolution 1. The frequency was calculated based on the total T cells, and a CLR transformation was applied to all samples.

Meskas, Justin, Sherrie Wang, and Ryan Brinkman. 2020. “FlowCut — An R Package for Precise and Accurate Automated Removal of Outlier Events and Flagging of Files Based on Time versus Fluorescence Analysis.” *BioRxiv*, June, 2020.04.23.058545.

Van Gassen, Sofie, Britt Callebaut, Mary J. Van Helden, Bart N. Lambrecht, Piet Demeester, Tom Dhaene, and Yvan Saeys. 2015. “FlowSOM: Using Self-Organizing Maps for Visualization and Interpretation of Cytometry Data.” *Cytometry. Part A: The Journal of the International Society for Analytical Cytology* 87 (7): 636–45.
